# Splicing accuracy varies across human introns, tissues and age

**DOI:** 10.1101/2023.03.29.534370

**Authors:** S García-Ruiz, D Zhang, E K Gustavsson, G Rocamora-Perez, M Grant-Peters, A Fairbrother-Browne, R H Reynolds, J W Brenton, A L Gil-Martínez, Z Chen, D C Rio, J A Botia, S Guelfi, L Collado-Torres, M Ryten

## Abstract

Alternative splicing impacts most multi-exonic human genes. Inaccuracies during this process may have an important role in ageing and disease. Here, we investigated mis-splicing using RNA-sequencing data from ~14K control samples and 42 human body sites, focusing on split reads partially mapping to known transcripts in annotation. We show that mis-splicing occurs at different rates across introns and tissues and that these splicing inaccuracies are primarily affected by the abundance of core components of the spliceosome assembly and its regulators. Using publicly available data on short-hairpin RNA-knockdowns of numerous spliceosomal components and related regulators, we found support for the importance of RNA-binding proteins in mis-splicing. We also demonstrated that age is positively correlated with mis-splicing, and it affects genes implicated in neurodegenerative diseases. This in-depth characterisation of mis-splicing can have important implications for our understanding of the role of splicing inaccuracies in human disease and the interpretation of long-read RNA-sequencing data.

## Introduction

RNA splicing is a crucial post-transcriptional process in which introns are excised from messenger RNA precursors (pre-mRNA), and exons are joined together to form mature mRNAs. It was previously believed that splicing of exons occurred in the order they appear in a gene. However, ~95% of human genes undergo alternative splicing (AS) whereby certain exons are differentially skipped resulting in different combinations of mature mRNA structures^1–3^. AS occurs within the nuclei of eukaryotic cells through base pairing between small nuclear ribonucleoproteins (snRNP) that form the spliceosome and the sequences signalling the intron boundaries, termed splice sites^4,5^. Splice site (ss) choice is largely regulated by splicing regulatory elements found throughout the pre-mRNA sequence^6–9^. Different RNA-binding proteins (RBPs) are then responsible for interacting with these regulatory signals to enhance or silence the recognition of introns and so activate or repress intron splicing accordingly within specific cells and tissues.

Splicing is a complex process and consequently accurate excision of an intron relies on overcoming multiple challenges. Firstly, the function of the spliceosome depends on the recognition and processing of a minimum of approximately 25 base pairs (bp) to correctly excise non-coding intronic sequences. This sets a relatively large mutational target in which germline and somatic variants could appear, compromising the correct identification of exon-intron boundaries^10–13^. Secondly, since some intronic sequences can be long (reaching lengths above 1 Mb^14^ in humans), cryptic splicing sequences will commonly exist within^15^, increasing the risk of decoy splice sites for spliceosome selection. Lastly, as observable in all biological systems, this process is subject to some stochastic variation^16–20^.

Accurate splicing is essential for human health^21–27^. While mechanisms such as the nonsense-mediated decay (NMD) can mitigate the impact of spurious mRNA transcripts^28–33^, differential use of splice sites escaping this mechanisms has demonstrated widespread dysregulation in a range of diseases^34^, including Alzheimer’s disease^35,36^, frontotemporal dementia^37,38^ and multiple cancers^39–42^. It has recently been suggested that ageing may exacerbate splicing errors, with intron retention events and spurious splicing becoming more prevalent with age and disease incidence^43^.

However, to the best of our knowledge, no study thus far has made a systematic attempt to evaluate the precision of splicing across introns, tissues, and age, while also modelling the various factors that could potentially regulate these processes. To address these questions, we used RNA-sequencing data provided by both the Genotype-Tissue Expression (GTEx) v8^44^ and the ENCODE consortium, and studied the accuracy in splicing of >300K annotated introns. We characterised mis-splicing and found robust patterns in its distribution reflecting the molecular architecture of spliceosome assembly and action. We identified local sequence conservation at splice sites as the most important and variable predictor of mis-splicing across tissues, which led us to investigate the role of RBP expression in tuning noise and changing its distribution. Given that RBP expression levels are already known to change across an individual’s lifespan, we studied the effect of age on mis-splicing. We demonstrated that age is positively correlated with mis-splicing and, in the human cortex, age-related increases in mis-splicing disproportionately affect genes implicated in neurodegenerative diseases. The analyses performed are summarised in **Fig. 1**. Altogether, these results show that mis-splicing is detectable across human tissues and modelling its characteristics provides novel insights into age-related human diseases.

**Fig. 1.**
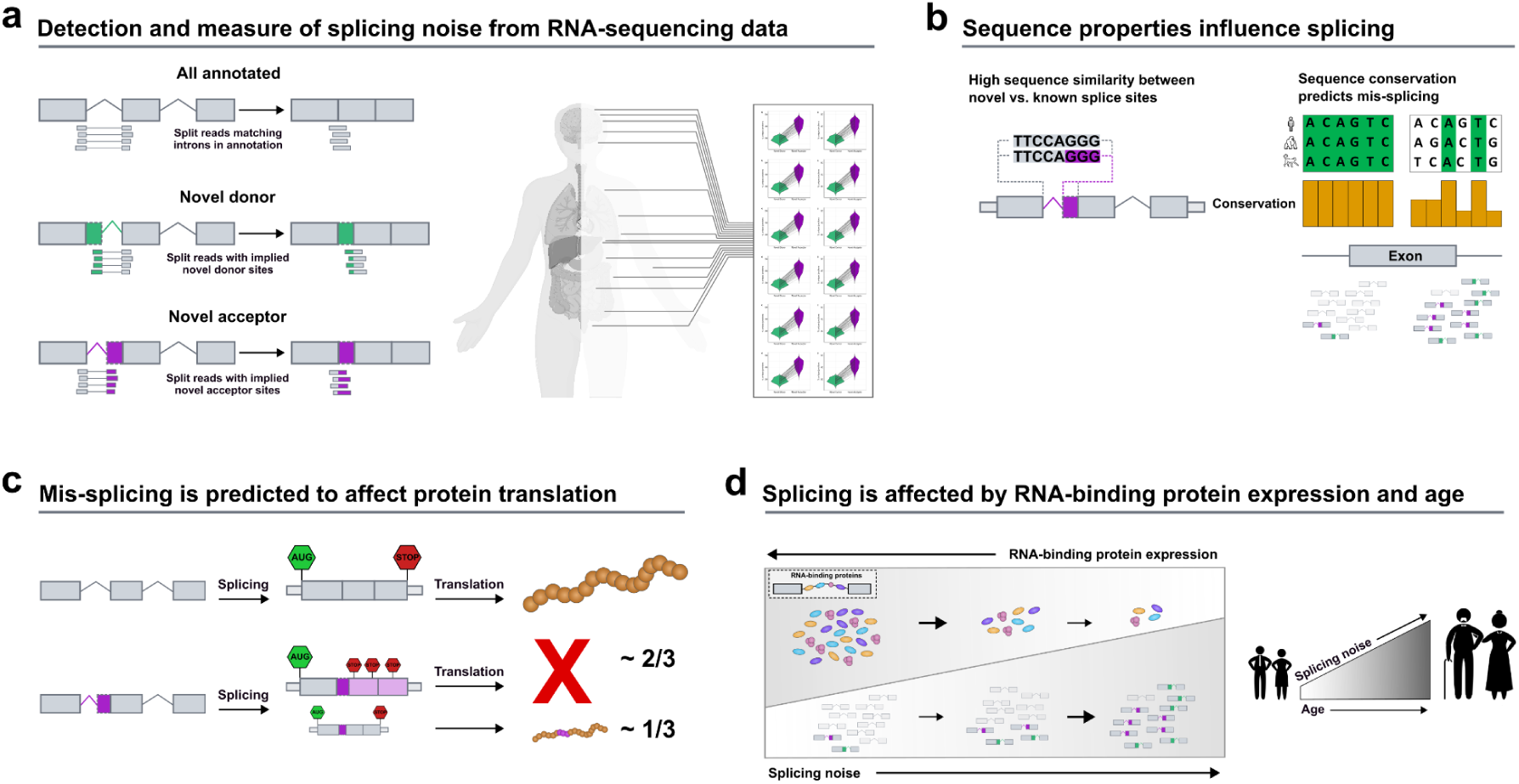
Overview of the analyses performed in this study. **a.** We studied splicing through three classes of split reads spanning exon-exon junctions: annotated, novel donor and novel acceptor split reads. The RNA-sequencing dataset used originated from the Genotype-Tissue Expression Consortium v8. In all 42 GTEx tissues studied, junctions from the novel acceptor category exceeded the number of unique novel donor junctions. **b.** Novel splice sites from the novel donor and novel acceptor junctions present high sequence similarity to annotated splice sites. Variability in mis-splicing rates across tissues is highly affected by inter-species sequence conservation at exon-intron boundaries. **c.** Novel junctions associated with protein-coding transcripts are predicted to be deleterious in 2/3 of cases. **d.** Reduced expression levels of the RNA-binding proteins (RBPs) responsible for sequence recognition appear to change splice site selection, which reduces the overall accuracy of the splicing process and increases mis-splicing rates. Age is positively correlated with mis-splicing increases in multiple human tissues.

## Results

### Novel donor and acceptor junctions are commonly detected and exceed the number of unique annotated introns by an average of 11-fold

Splicing events can be accurately detected from short-read RNA-sequencing data using split reads. Split reads are reads that map to the genome with a gapped alignment, indicating the excision of an intron. We focused on three classes of split reads: i) annotated exon-exon junction reads, which precisely match an intron within annotation (Ensembl-v105), ii) novel donor junctions, where only the implied acceptor site matches an intron-exon boundary within annotation, and iii) novel acceptor junctions, where only the implied donor site matches an exon-intron boundary within the annotation (**Fig. 1**). To study splicing through these three junction classes, we leveraged RNA-sequencing data processed by the relational database, IntroVerse^45^, and originating from the Genotype-Tissue Expression Consortium^44^ (GTEx) v8 data set. Using a subset of the data provided by IntroVerse relating to 324,956 annotated introns (**Extended Data Fig. 1a,b**), we studied all their linked novel donor and acceptor junctions. We found that 268,988 (82.8%) annotated introns had at least one associated novel junction, with only 55,968 annotated introns appearing to be accurately spliced across all samples and 42 tissues studied. Collectively, we detected 3,865,268 unique novel donor and acceptor junctions, equating to 14 novel junctions per annotated intron. The detection of unique novel donor and acceptor junctions was a common finding across all tissues, with the highest numbers found in *“Cells - EBV-transformed lymphocytes”* tissue and the lowest in *“Whole Blood”*.

### Over 98% of novel donor and acceptor junctions are likely to be generated through splicing errors

Unique novel junctions may represent novel transcripts, but given the high numbers detected, novel junctions could also be the product of splicing errors. To explore this, we leveraged the existence of multiple reference Ensembl transcriptome builds, namely v97 (May, 2019) and v105 (June, 2021), assuming an increased accuracy over their 2-year gap. For each tissue, we re-processed and re-annotated each split read provided by the GTEx v8 to the v97 and v105 annotation builds **(Methods)**. We found that across all tissues, on average only 0.8% [range:0.55-1.23%] of junctions defined as novel donor or acceptor junctions using v97 were reclassified as annotated introns in v105, and thus part of a transcript structure (**Fig. 2a**). Interestingly, we noted that the highest re-classification rates were observed amongst human brain tissues, on average 0.8±0.12%. We predicted that the re-classification of novel junction reads should decrease with successive annotations and so we extended our analysis to frontal cortex brain tissue to include Ensembl versions published from 2014 to 2021. The reclassification rate of novel junctions in frontal cortex decreased incrementally from 2.36% to 0.33% **(Extended Data Fig. 2),** consistent with previous studies reporting that the number of novel junctions has been plateauing since 2013^46^. These findings suggest that the vast majority of novel junctions are generated through mis-splicing events, with on average <0.8% being explained by junctions originating from stable transcripts.

**Fig. 2.**
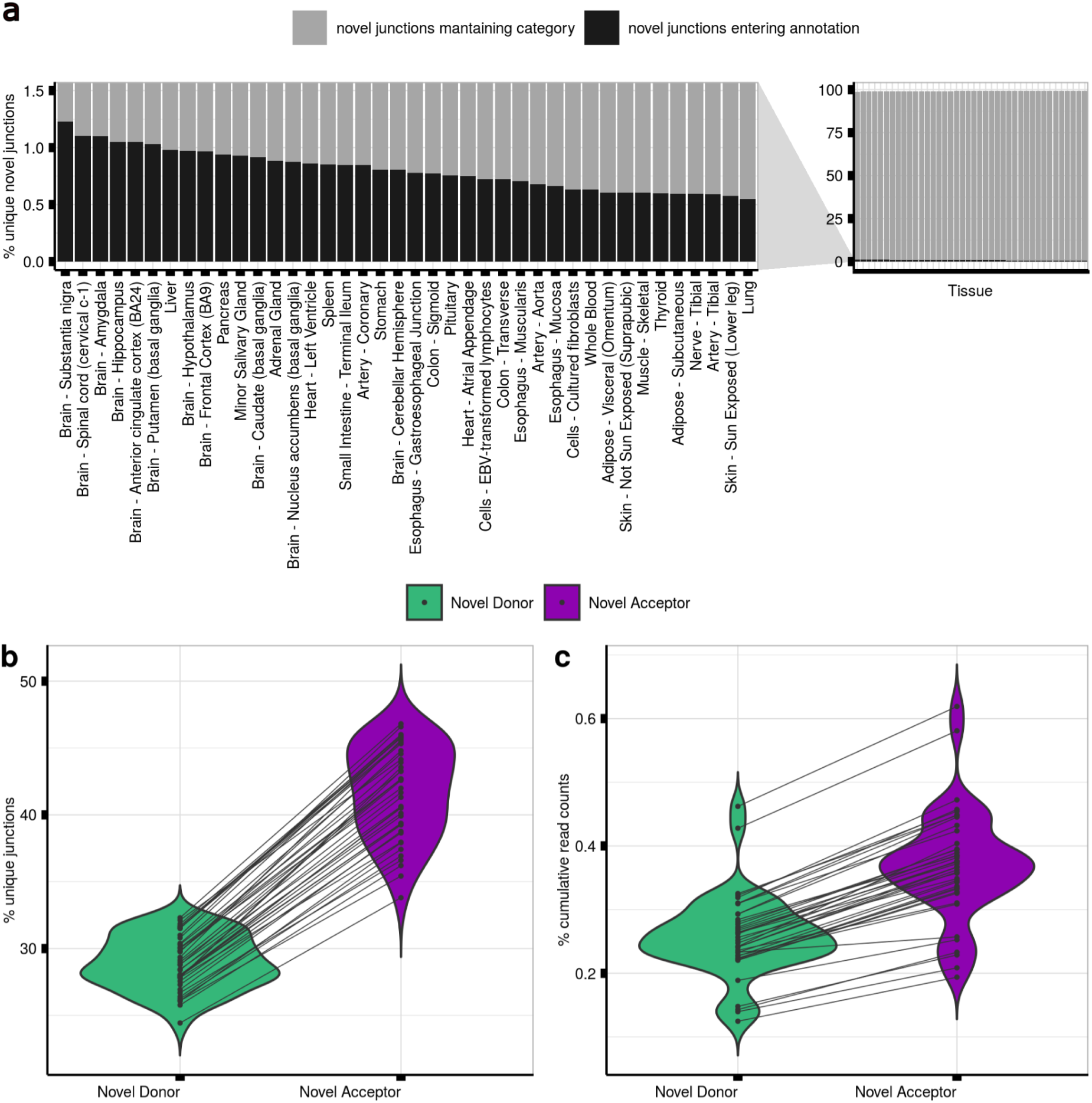
Mis-splicing can be measured using short-read RNA-sequencing data. **a.** Contamination rates in Ensembl v97 as compared to Ensembl v105 per GTEx tissue. Bars in dark grey represent the percentage of split reads classified as novel junctions using Ensembl v97 that entered annotation as annotated introns in Ensembl v105. Bars in light grey represent the percentage of novel junctions in Ensembl v97 that maintained novel category in Ensembl v105. **b.** Percentage of unique novel donor and novel acceptor junctions per GTEx tissue in Ensembl v105. The crossing lines link the percentage of novel donor and novel acceptor junctions found in each body site. **c.** Percentage of cumulative number of read counts that the novel donor and novel acceptor junctions present per GTEx tissue in Ensembl v105. The crossing lines link the percentage of cumulative number of novel donor and novel acceptor read counts found in each body site.

### Mis-splicing is more common at acceptor than donor splice sites

The recognition of the donor splice site (5’ss) and acceptor splice site (3’ss) of an intron is performed by separate components of the splicing machinery^34,47,48^. We aimed to test whether mis-splicing rates at these splice sites also differed. To assess this, we compared the numbers of unique novel donor and acceptor junctions detected in each tissue to the numbers of unique annotated introns. We found that novel donor and acceptor junctions consistently accounted for the majority of unique junctions detected (70.6% [range:58.2-79.1%]), and that the novel acceptor category exceeded the novel donor across the samples of all tissues (**Fig. 2b**). While we detected an average of 241,044 unique annotated introns across all tissues, unique novel donor and acceptor junctions averaged 249,366 and 360,400, respectively.

We reasoned that while mis-splicing might generate high numbers of unique novel junctions in a sample, each of these junctions would be expected to have a low number of associated reads. Consistent with this prediction, we found that novel donor and acceptor junctions together accounted for 0.32-1.08% of all junction reads whereas annotated introns accounted for 98.92-99.68% of the junction reads across all tissues evaluated (**Fig. 2c**). Focusing on frontal cortex brain tissue, we found that this equated to a median read count of 2,694 reads per annotated intron with novel donor and acceptor junctions having a median read count of only 2 reads in both cases. These findings were replicated across all human tissues (**Supplementary Table 1**) and were consistent with novel junctions generated through splicing errors.

### High sequence similarity between novel splice sites and their annotated pairs explains mis-splicing

Sequences delineating intron boundaries are diverse and cryptic splice sites have the potential to induce mis-splicing events when present in close proximity to them^49^. We applied the MaxEntScan^50^ (MES) algorithm to assess the similarity of all annotated and novel 5’ss and 3’ss to consensus representative sequences in humans. We found significant overlaps between the distribution of MES scores assigned to annotated versus novel splice sites, suggesting that the splicing machinery would be expected to recognise the latter (**Extended Data Fig. 3**).

Given that splice selection is likely to be a competitive process, we leveraged our paired data structure to compare MES scores between annotated and novel junction pairs (termed delta MES score). We found that the majority of novel 5’ss and 3’ss were weaker than the corresponding annotated site, with 82.6% of novel 5’ss and 85.8% of novel 3’ss having positive delta MES scores (**Fig. 3a,b**). Moreover, novel 5’ss and 3’ss had a median delta value of 3.6 and 5.2, respectively, in keeping with the higher number of novel acceptor as compared to novel donor junctions detected in all tissues, and similar MES scores to their annotated pairs. Overall, these results suggest that the strength of local splicing signals is not sufficient to guarantee accurate splicing^13,51^.

**Fig. 3.**
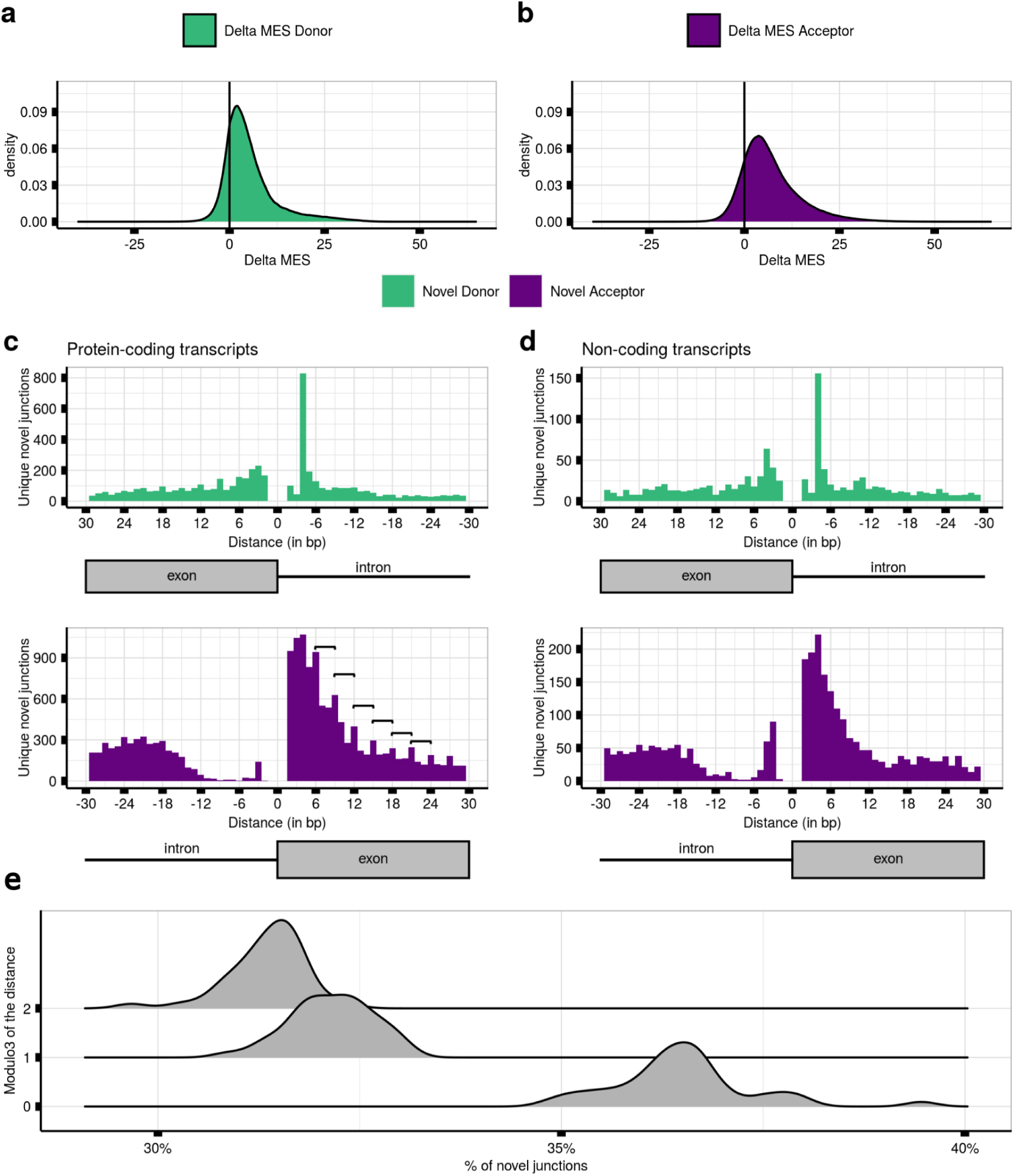
Mis-splicing is explained by the high sequence feature similarity between novel splice sites and their annotated pairs. **a.** MaxEntScan (MES) Delta scores between the scores assigned to the 9-bp sequence at the 5’ss of the annotated introns and their novel donor pairs across all tissues. **b.** MES Delta scores between the scores assigned to the 23-bp sequence located at the 3’ss of the annotated introns and their novel acceptor pairs across all tissues. **c.** Distances lying between the novel splice site of each novel junction and their annotated pairs from protein-coding transcripts in frontal cortex brain tissue. **d.** Distances lying between the novel splice site of each novel junction and their annotated pairs from non-protein-coding transcripts in frontal cortex brain tissue. **e.** Modulo3 of the distances between each novel junction and linked annotated intron to a maximum distance of 100 bp within MANE transcripts from all body sites.

### Novel junctions associated with protein-coding transcripts are predicted to be deleterious in 63.5% of cases

High sequence similarity between novel and annotated splice sites might be expected if these sites were near each other. Thus, we analysed the relationship between annotated and novel splice sites in close proximity, focusing on the distribution of the latter within 30 bp upstream and downstream of annotated sites in frontal cortex brain tissue (**Methods**). We noted that: i) both novel 5’ss and 3’ss were located near to paired annotated sites; ii) the distribution of mis-splicing was different between annotated 5’ss (mode=−4bp/3bp) and 3’ss (mode=−21bp/4bp); and iii) mis-splicing was highly asymmetric at annotated acceptor sites, with a very low mis-splicing density upstream this intron-exon boundary, suggesting that this mis-splicing pattern was driven by the AG exclusion zone (AGEZ). These results were replicated across all tissues (**Supplementary Table 2**), consistent with novel junctions originating from splicing errors.

We also observed regular splice site peaks occurring at 3 bp intervals, most apparent in novel acceptor events downstream of the paired annotated site, namely within annotated exons. Using data from frontal cortex brain tissue, we noticed that these peaks were only observed in protein-coding transcripts (n=35,654) (**Fig. 3c,d**), suggesting that they could be generated by the preferential degradation of deleterious transcripts through NMD.

To further explore this possibility, we studied the divisibility by 3, equating to the size of a codon, of the distances between each novel junction and their linked annotated 5’ss and 3’ss. Focusing on splice sites exclusively used in protein-coding transcripts in frontal cortex **(Methods)**, this analysis demonstrated that 62.5% of all novel sites were located at distances not divisible by 3, implying that these splicing events would result in a deleterious frameshifts for downstream translation events. When focusing on each modulo3 value independently, we observed an overall preference to maintain codon reading frame (mod3=0, 37.4%; mod3=1, 31.4%; mod3=2, 31.2%). Across all tissues, 63.55% of the novel junctions would likely disrupt the reading frame **(Fig. 3e),** supporting the view of novel junctions originating from splicing errors.

### Mis-splicing rates vary across introns and are likely to be underestimated in bulk RNA-seq data

Next, we wondered if splicing fidelity varies across introns and genes across the genome. We used the Mis-Splicing Ratio measures to assess the frequency of mis-splicing at both the 5’ss (*MSR_D_*) and 3’ss (*MSR_A_*) of each annotated intron (**Methods**). Focusing on frontal cortex brain tissue, we observed that while splicing errors are detected infrequently, with the *MSR_D_* and *MSR_A_* values highly skewed towards low values, there was considerable variation across introns (*MSR_D_*IQR=7.2e-04; *MSR_A_*IQR=1.9e-03). Furthermore, consistent with the overall higher detection of novel acceptor as compared to novel donor junctions, we observed a significant difference in the two *MSR_D_* and *MSR_A_* distributions (V=8e7, pval<2.2e-16) **(Fig. 4a)**.

**Fig. 4.**
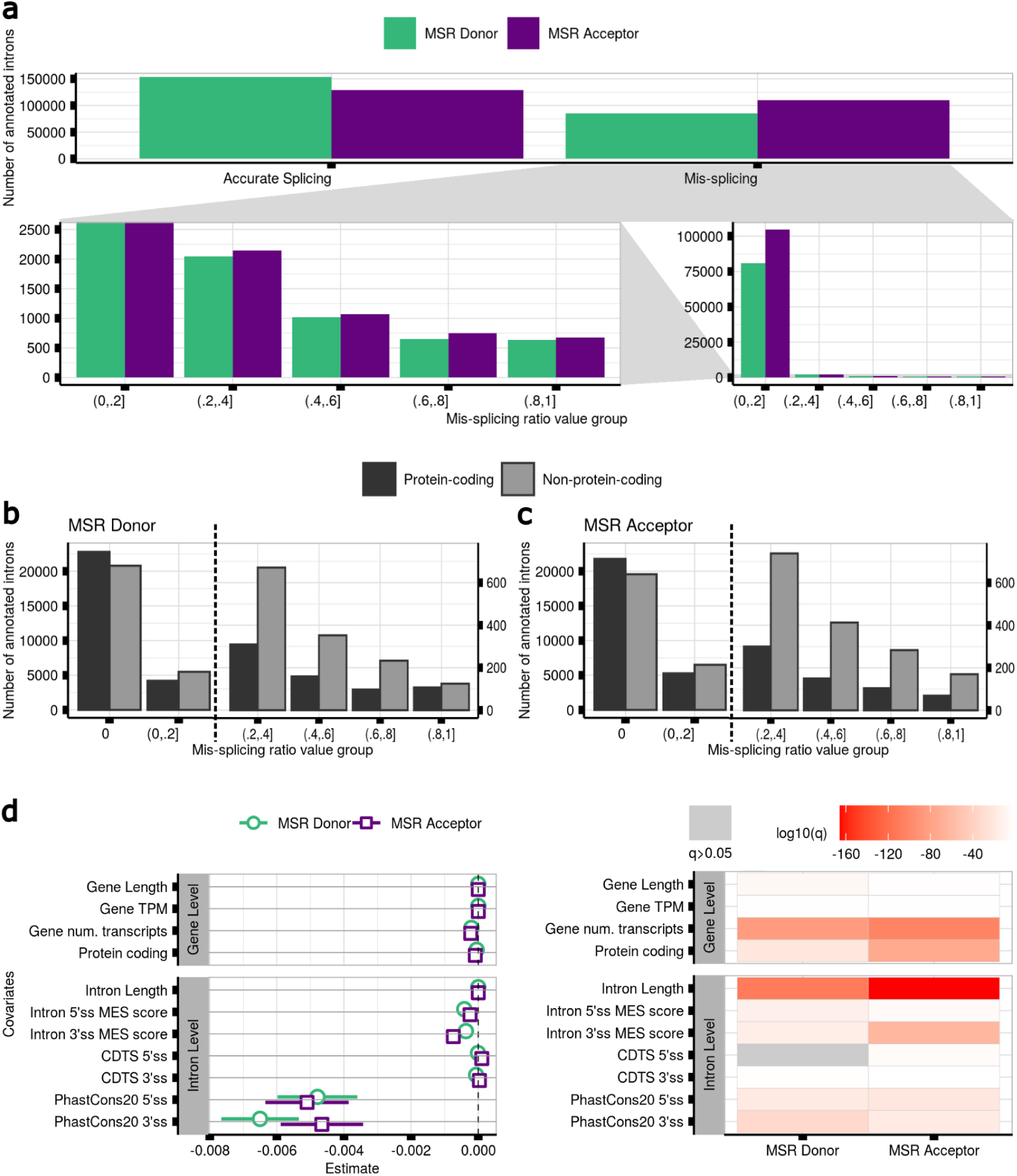
Mis-splicing rates vary across introns and local sequence conservation is its most important predictor. **a.** Mis-splicing rates occurring at the 5’ss and 3’ss of the annotated introns in samples of frontal cortex brain tissue. Bottom right: mis-splicing rates for mis-spliced introns across binned values. Bottom left: a zoomed in view of the bottom right panel, with the y-axis cropped. **b.** Mis-splicing rates occurring at the 5’ss of the annotated introns located within protein-coding vs non-protein-coding transcripts in samples from frontal cortex brain tissue. The black dashed vertical line separates the bars represented under the two different y-scales displayed. Right y-scale: a zoomed in view of the y-axis scale on the left side. **c.** Mis-splicing rates at the 3’ss of the annotated introns located within protein-coding vs non-protein-coding transcripts in samples from frontal cortex brain tissue. **d.** Linear regression models to predict the mis-splicing rates at the 5’ss and 3’ss of the annotated introns in samples from frontal cortex brain tissue. P-values were corrected for multiple testing using the Benjamini-Hochberg method, resulting in q-values. Grey values represent non-significant q-values.

Given that NMD activity would be expected to reduce the detection of splicing errors amongst mRNA transcripts, we compared mis-splicing of annotated introns in protein-coding versus non-coding transcripts **(Extended Data Fig. 4) (Methods)**. We found that mis-splicing is more frequent amongst annotated introns from non-protein-coding transcripts at both the 5’ss (V=1.3e7,pval<2.2e-16) and 3’ss (V=1.5e7, pval<2.2e-16) **(Fig. 4b,c)**, suggesting that the frequency of splicing errors is likely to be under-estimated. These findings were validated across all tissues (**Supplementary Table 3**).

### Local sequence conservation is the most important predictor of mis-splicing

Given the variability in mis-splicing across introns, we wanted to identify features that could influence its generation. We therefore built two linear regression models to predict the rate of mis-splicing as defined by *MSR_D_* and *MSR_A_* values, and used as predictors different features of each annotated intron and the gene in which it was located **(Methods)**. This analysis yielded three main findings. Firstly, we found that gene-level features had a small but significant effect on mis-splicing at both splice sites. Increases in gene length, expression levels and associated transcript number predicted a reduction in mis-splicing, suggesting that splicing inaccuracies within highly expressed genes and high transcript diversity might be energetically costly for organisms^20,52^ and so selected against (**Fig. 4d**). Secondly, this analysis provided support for splice site inter-communication^34,53^, with sequence properties at the 3’ss impacting on the fidelity of 5’ss splicing and to a lesser extent vice versa. Finally, we found that sequence conservation in genomic regions flanking the 5’ and 3’ss had the largest effect on splicing accuracy, with higher conservation scores at both sites associated with lower mis-splicing. Interestingly, the estimate values for sequence conservation at both splice sites were much larger (*MSR_D_*=[-4.8e-03,-6.5e-03], *MSR_A_*=[-5.1e-03,-4.6e-03]) than those associated with CDTS scores (*MSR_D_*=[-6.5e-06,-6.3e-05], *MSR_A_*=[1.1e-04,3.5e-05]). Given that the latter is a measure of sequence constraint amongst humans, this would suggest that germline genetic variation across individuals is not a major driver of mis-splicing.

### Accuracy in splicing is affected by RNA-binding protein (RBP) expression changes

To better understand the factors influencing mis-splicing across tissues, we expanded our analyses to all tissues, focusing on a common set of annotated introns (n=151,729) **(Methods)**. We found that gene-level properties were significantly associated with mis-splicing in all tissues and that the effect of sequence conservation on splicing inaccuracies varied at both splice sites (*MSR_D_*=[-7.5e-03,-3.2e-03] **(Fig. 5a)**, *MSR_A_*=[-6.6e-03,-3.7e-03] (**Fig. 5b**)), despite the conservation scores being the same across tissues. We hypothesised that somatic mutation rates in tissues might affect critical splicing sequences, causing changes in their recognition by RBPs and resulting in splicing errors. However, we did not find any significant differences in mis-splicing across annotated introns between *“Skin - Sun Exposed”* and *“Skin - Not Sun Exposed”* (*MSR_D_*V=5.2e9, pval=3.9e-01, and *MSR_A_* V=5.9e9, pval=9.3e-01) (**Methods**, **Extended Data Fig. 5**).

**Fig. 5.**
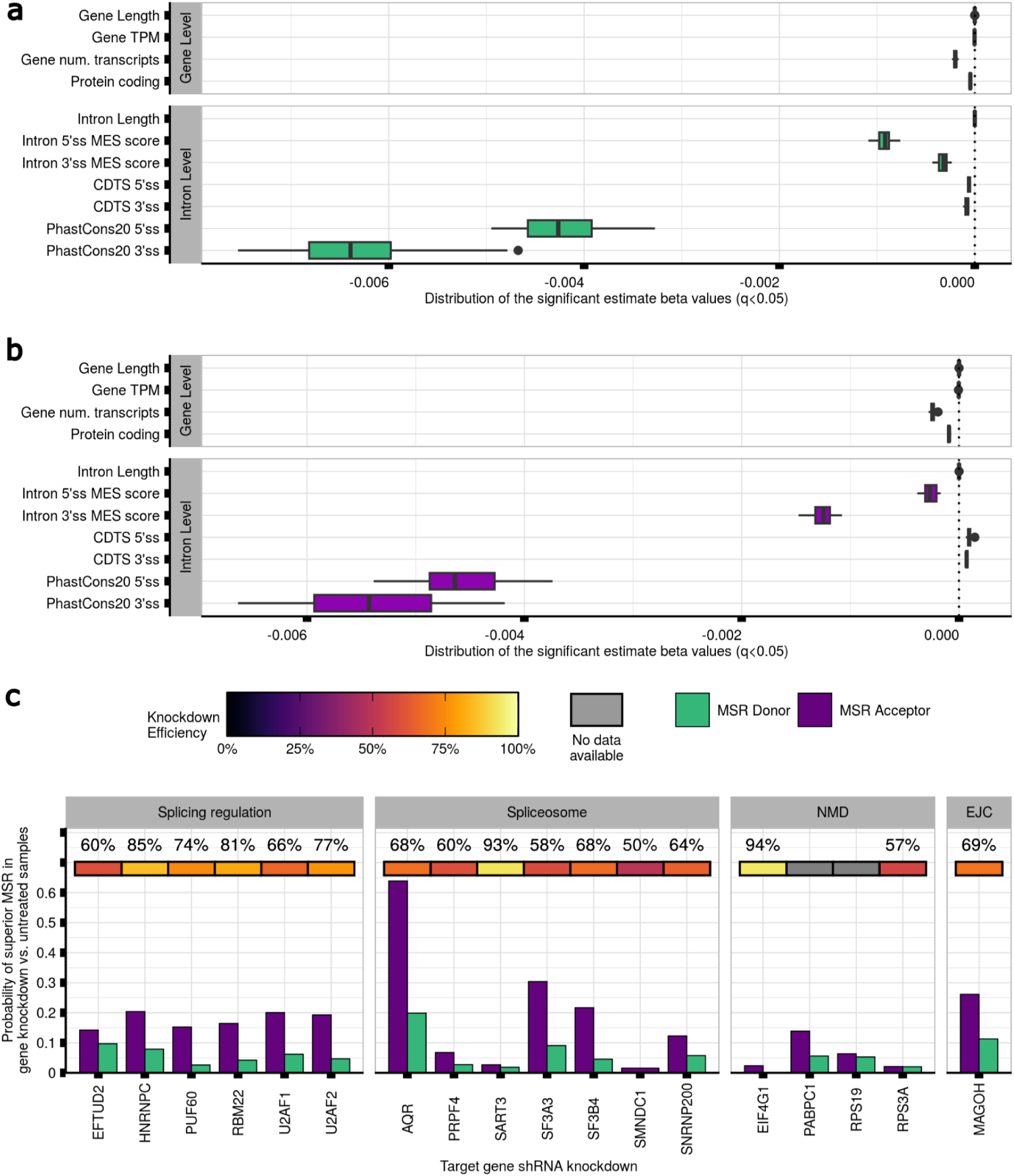
Mis-splicing rates vary across tissues and it could be explained by variable RNA-binding protein (RBP) expression. **a.** Prediction of the mis-splicing rates at the 5’ss of the introns studied across 42 body sites. Every boxplot contains 42 beta estimate values. **b.** Prediction of the mis-splicing rates at the 3’ss of the introns studied across 42 body sites. Every boxplot contains 42 beta estimate values. **c.** Probability of superior mis-splicing rates at the 5’ss and 3’ss of the annotated introns in samples with the shRNA knockdown of each gene as compared to untreated samples. The top heatmap track contains the knockdown efficiency of the associated protein. Grey values represent no available data for the knockdown efficiency of the associated protein.

Based on these findings, we considered if tissue-specific expression levels of RBPs involved in splicing processes could explain the effect of sequence conservation on mis-splicing (**Methods**, **Extended Data Fig. 6, Supplementary Fig. 1,2**). To explore this possibility, we analysed ENCODE data involving knockdowns of 56 genes related to splicing regulation, spliceosome assembly, exon-junction complex recognition^54^ and NMD. Our analysis revealed a significant increase in mis-splicing rates in samples with gene knockdowns compared to untreated controls for 89.3% (*MSR_D_* q<7.4e-50,n=50) and 91.1% (*MSR_A_* q<1.2e-03,n=51) of the 56 genes considered. Knockdowns of the splicing machinery components tended to have a greater effect on 3’ss than 5’ss mis-splicing (mean *MSR_D_*effsize=0.09 [0.02,0.37]; mean *MSR_A_*effsize=0.1 [0.01,0.62]), except for 6 genes (**Supplementary Table 4, 5**). Notably, AQR, EIF4A3, SF3A3, U2AF1, U2AF2, and MAGOH knockdowns resulted in the highest increases in 5’ss and 3’ss mis-splicing (**Fig. 5c**).

Our analysis also revealed distinct patterns in mis-splicing distribution depending on the gene targeted. For instance, knocking down U2AF2 expression led to a significant increase in mis-splicing within 15-30 bp upstream of the annotated acceptor site. Delta MES values indicated weaker 3’ss were being targeted for the novel acceptor junctions (W=2.62e+08, pval<9.4e-04), suggesting that the spliceosome was no longer able to accurately distinguish splicing signals at acceptor sites **(Fig. 6a, b)**. Similarly, AQR knockdowns resulted in a remarkably high number of mis-splicing within the 15-200 bp sequence window upstream of the annotated 3’ss **(Fig. 6c)**, with a further reduction in the strength of novel 3’ss compared to paired annotated sites (W=2.2e-09, pval<2.2e-16) **(Fig. 6b)**.

**Fig. 6.**
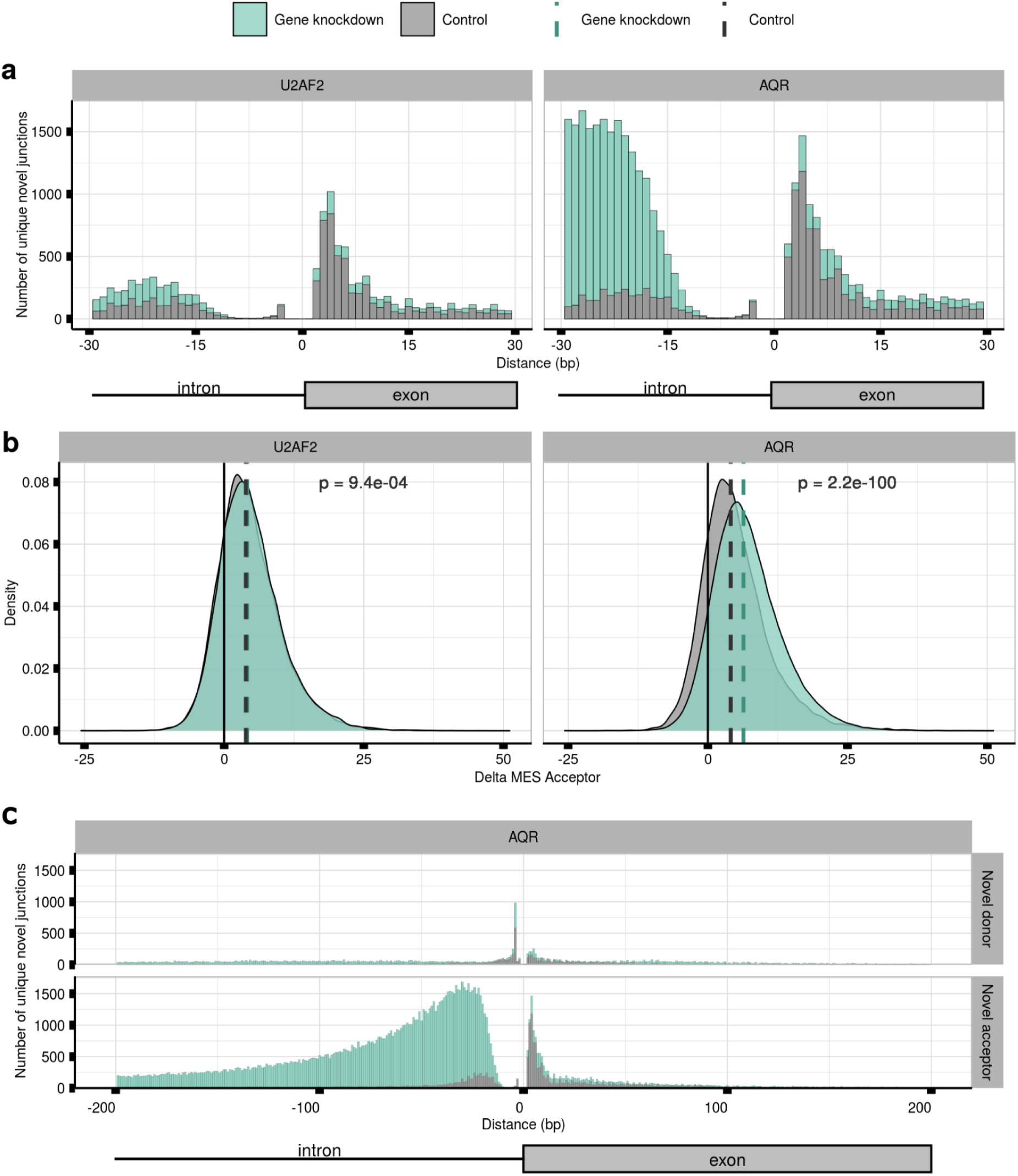
shRNA knockdown of U2AF2 and AQR produces different patterns of mis-splicing distribution. **a.** Distances lying between the novel 3’ss of each novel acceptor junction and their annotated pairs in samples with the shRNA knockdown of the U2AF2 and AQR genes, respectively, as compared to untreated controls. **b.** MES Delta scores between the scores assigned to the 23-bp sequence located at the 3’ss of the annotated introns and their novel acceptor pairs in samples with the shRNA knockdown of the U2AF2 and AQR genes, respectively, as compared to untreated controls. Dashed vertical lines represent the median value of each distribution. **c.** Distances lying between the novel 3’ss of each novel acceptor junction and their annotated pairs in samples with the shRNA knockdown of the AQR gene and compared to untreated samples up to a distance of 200 bp.

### Increasing age is associated with increasing levels of mis-splicing

Previous studies have reported an overall reduction in the expression of multiple RBPs with increasing age^55–61^. We formally assessed this in the GTEx data set and, focusing on brain tissue, found that the expression levels of 20.7% of the 116 RBPs studied were negatively affected by age (1.1e-07<q<4.2e-02) **(Methods**, **Supplementary Table 6)**. To investigate if age-related changes in RBP expression could result in increasing levels of mis-splicing, we grouped samples for each body site into 2 extreme age clusters, “20-39” and “60-79” years and, after controlling for potential confounding covariates, we selected a set of n=206,067 annotated introns shared across age groups and body sites **(Methods, Extended Data Fig. 7,8)**. We found that *MSR_D_* values in the “60-79” age group were significantly higher than those in the “20-39” cluster in 9 of the 18 body sites analysed (4.1e-02<effsize<0.1; 2.22e-100<q<1.01e-43). Similarly, *MSR_A_* values in the “60-79” age group were significantly higher than those in the “20-39” category in 11 of the 18 tissues assessed (8.1e-03<effsize<1.4e-01; 2.22e-100<q<1.14e-02). In both cases, the highest effect size was found in blood vessel tissue (**Fig. 7a, Supplementary Table 7**).

**Fig. 7.**
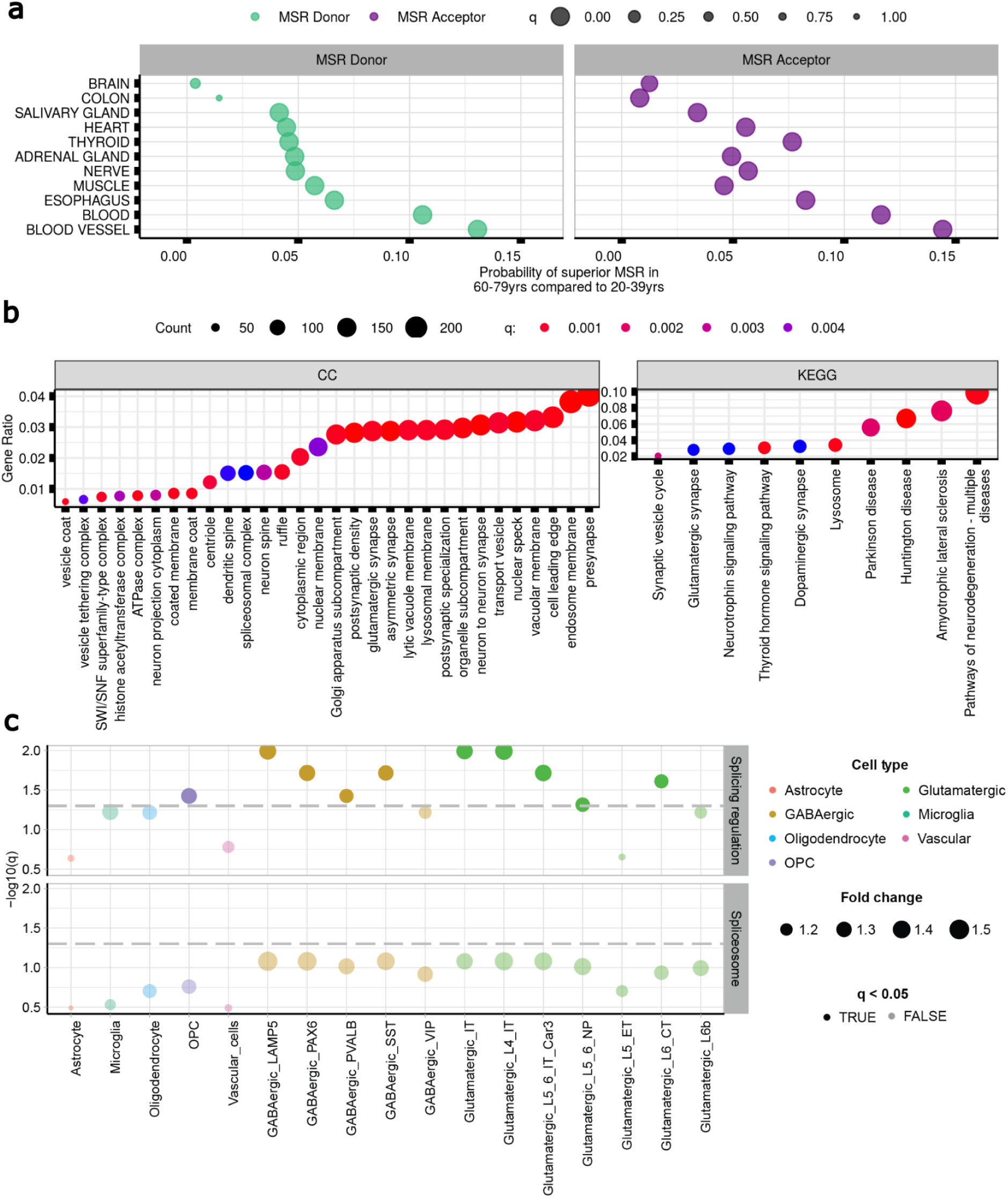
Mis-splicing increases with age and affects genes involved in neuronal function. **a.** Probability of superior mis-splicing rates at the 5’ss and 3’ss of the annotated introns in samples from donors aged between 60-79 years-old as compared to 20-39 yrs. **b.** GO and KEGG enrichment analysis of the genes containing introns with increasing levels of mis-splicing rates with age (i.e. 20-39yrs < 60-79yrs) at their 5’ss and/or 3’ss in samples from brain tissues. **c.** Cell-type specific expression of 111 splicing-regulator and spliceosomal RBPs (Van Nostrand et al. 2020) in cell types derived from multiple cortical regions of the human brain (Shen et al. 2012). The cell type annotations used correspond to 7 subtypes of glutamatergic neurons, 5 subtypes of GABAergic neurons, Astrocyte, Microglia, Oligodendrocyte, OPC, and vascular cell (Endothelial, Pericyte and, VLMC) as non-neuronal cell types. The dashed grey horizontal lines represent the minimum level of significance, with dots displayed above the line showing significant specific expression for a given cell type. P-values were corrected for multiple testing using the Benjamini-Hochberg method, resulting in q-values.

Given the complexity of splicing in the human brain and importance of age-related disorders affecting this organ, we further investigated the properties of introns with evidence of age-related increases in mis-splicing in brain. We identified n=14,714 annotated introns of interest based on their *MSR_D_* or *MSR_A_* increasing values (**Methods**). After assigning these introns to their unique genes n=8,117, we used Gene Ontology (GO) Enrichment analysis to determine if age-related increases in mis-splicing might have an impact on specific biological processes or pathways. Interestingly, this analysis identified significant enrichment in terms such as *“neuron to neuron synapse”* (q<2.3e-06), *“tau protein binding”* (q<4.8e-04) and *“dendritic spine”* (q<4e-04) (**Fig. 7b, Supplementary Table 8,9**). Since the former term suggested that mis-splicing might affect neurons more than other cell types, we assessed cell-type specific expression of RBPs in the human brain. Using single-nucleus RNA-sequencing data from the Allen Brain Atlas covering multiple cortical regions^62^, we investigated the cell-type specificity of 111 splicing-regulator and spliceosomal RBPs^54^ across all major cell types. We found that splicing-regulator RBPs were more highly expressed than would be expected by chance in the following cell-types: oligodendrocyte precursor cells (OPCs), 4 subtypes of GABAergive neuron and 5 subtypes of glutamatergic neuron **(Fig. 7c, Extended Data Fig. 9, Supplementary Table 10,11)**. The enrichment of RBPs within specific neuronal cell types suggests that neurons may be particularly sensitive to changes in RBP expression, and by extension, particularly vulnerable to age-related increases in mis-splicing.

## Discussion

Here we have shown that mis-splicing events are common across human tissues and occur near annotated intron-exon boundaries in a distinctive and predictable manner. Mis-splicing rates are higher at acceptor sites than at donor sites, and non-coding transcripts have higher mis-splicing rates at both sites in all tissues. We discovered that mis-splicing rates vary across introns and tissues, and are predictable based largely on local sequence properties. Reduced expression of spliceosome components and regulators is a significant contributing factor to the variability in mis-splicing, as evidenced by *in vitro* knockdowns of RBPs and *in vivo* with ageing. Additionally, in the ageing human brain, mis-splicing disproportionately affects genes involved in neuronal function and proteostasis, with implications for age-related neurodegenerative disorders.

One of the most striking and robust findings in this study was the consistently higher accuracy of 5’ss as compared to 3’ss recognition. This is likely to reflect intrinsic weaknesses and molecular differences in these processes. Initial recognition of the 5′ss of an intron is carried out by the U1 snRNP complex of the spliceosome. Even though their base-pairing interactions are often imperfect, this process is thought to be highly efficient^48,63,64^. In contrast, recognition of the 3’ end of introns requires cooperative binding of three interacting proteins to three neighbouring sequence motifs. Besides, a given 3’ss can be associated with more than one functional branch point^65^. Our findings support this view and suggest that this complexity makes this process particularly sensitive to errors.

There are a range of ways in which splicing errors could arise at both splice sites. Most simply, they could originate from genomic sequence variation due to germline and somatic mutations or inaccuracies in the recognition of splicing signals by the spliceosome machinery itself. We found limited evidence to support the former. Using a measure of DNA sequence constraint in humans, namely CDTS scores^66^, we found that while germline genetic variation at exon-intron boundaries influenced mis-splicing rates, the effect was consistently small across tissues. Similarly, when we assessed the potential impact of somatic mutations by comparing mis-splicing rates in unexposed versus sun-exposed skin (known to have a higher somatic mutation load^67^), we found no significant differences. In fact, local sequence conservation across species had the highest effect on mis-splicing across tissues, despite conservation scores across exon-intron boundaries being the same in all tissues.

These findings are consistent with the current understanding of splicing and its evolution. While splicing is thought to have arisen through the self-removal of introns from primitive RNA molecules^68^, it is postulated that their strict sequence and structural requirements progressively relaxed over time^69^. Consequently, these introns became more reliant on accurate expression of spliceosome RNAs and proteins for efficient recognition of cis-intronic splicing sequences and proper splicing. We suspected that the variable effect of sequence conservation on mis-splicing across human tissues could be explained by differences in the expression of these components, making mis-splicing primarily a problem of inaccurate sequence recognition.

We formally assessed this hypothesis using publicly available data from the ENCODE consortium to measure mis-splicing rates following shRNA knockdown of multiple RBPs^54^. Interestingly, we found that depending on the RBP targeted, there were distinctive patterns of mis-splicing distribution, suggesting a dependency on adequate levels of expression of each spliceosomal component to accurately target a splice site. Surprisingly, shRNA knockdowns of core spliceosomal molecules, such as the U2AF2 and AQR, did not reduce the total levels of splicing activity. Instead, these knockdowns appeared to change splice site selection, reducing the overall accuracy of this process. Certainly, mutations in the U2AF heterodimer have been found to be rate-limiting for splice site choice^70–72^.

Given that changes in the activity of core spliceosomal components have also been linked to a broad range of pathologies^73–76^ and ageing^25,60,77,78^, we studied mis-splicing changes with age in a range of tissues. This analysis revealed an increase in mis-splicing in the eldest group across most body sites. Focusing on human brain due to the known importance of RBPs in brain diseases^75,76^, we noted that core spliceosomal genes and genes involved in synaptic function and proteostasis were disproportionately affected by age-related changes in mis-splicing. This could be due to higher requirements for RBP expression in neurons, as suggested by our cell-type specificity analysis.

We note that our findings on mis-splicing rates also have the potential to inform long-read RNA-sequencing analyses which have tended to identify large numbers of novel transcripts particularly when performed at high depth^79^. More specifically, an awareness of patterns in splicing noise could help differentiate between low-expressed novel isoforms of process biological interest and transcriptional noise. Besides, the success in splicing-modulating RNA therapies in muscular dystrophy and cancer cells^80–82^, elucidates the potential that a deeper understanding of splicing and its regulatory mechanisms can have to help design new therapies that counteract the detrimental effects of these pathologies.

However, we note some important limitations of this study. Firstly, all analyses have been performed using bulk RNA-sequencing data despite our own analyses of RBP expression by cell type. This is likely to impact on our assessment of mis-splicing rates and its biological impact, potentially leading to an underestimate of its effect on rarer cell types. Furthermore, our analyses have not attempted to model the impact of the NMD. Given the importance of this pathway for the degradation of potentially deleterious novel transcripts, we postulate that the mis-splicing events we observe are primarily those that have partially escaped NMD suggesting that mis-splicing rates are in fact higher across all human tissues.

Nonetheless, taken together, our results show that mis-splicing is common and that understanding its patterns will inform our understanding of the role of splicing in senescence, healthy ageing and disease. We believe that this will be key to the successful application of RNA-targeting therapies particularly in human brain.

## Online Methods

### RNA-sequencing data download and processing

We downloaded and processed data from the IntroVerse database^45^, which contains the splicing activity of 332,571 annotated introns (as defined by Ensembl-v105) and a linked set of 1,950,821 novel donor and 2,728,653 novel acceptor junctions, covering 17,510 human control RNA samples and 54 tissues. This dataset of exon-exon junctions was originally provided by the Genotype-Tissue Expression Consortium (GTEx) v8^44^ and processed by the recount3^83^ (version 1.0.7, https://github.com/LieberInstitute/recount3) project.

The Illumina TruSeq library construction protocol (non-stranded 76 bp-long reads, polyA+ selection) was used in GTEx v8. Samples from GTEx v8 were processed by the recount3 project through Monorail^83^ (version 1.0.0, https://github.com/langmead-lab/monorail-external) which uses STAR^84^ (RRID:SCR_004463, http://code.google.com/p/rna-star/) to detect and summarise exon-exon splice junctions for each sample. Megadepth^85^ (version 1.0.3, RRID:SCR_022779, https://github.com/ChristopherWilks/megadepth) was also used by recount3 to analyse the BAM files output by STAR (version 2.7.3a, RRID:SCR_004463, http://code.google.com/p/rna-star/).

IntroVerse uses the Bioconductor R package dasper^86^ (version 1.4.3, http://www.bioconductor.org/packages/dasper) to annotate the split reads (Ensembl-v105) from GTEx v8 and processed by recount3. Within IntroVerse each novel donor and acceptor junction is first carefully quality-controlled (to ensure that novel junctions could feasibly arise through splicing) and then assigned uniquely to a specific annotated intron. Among the quality-control criteria applied by IntroVerse, all split reads shorter than 25 base pairs (bp) were discarded as well as all split reads located within unplaced sequences on the reference chromosomes and overlapping any of the regions published within the hg38 ENCODE Blacklist^87^ (v2.0, https://github.com/Boyle-Lab/Blacklist/blob/master/lists/hg38-blacklist.v2.bed.gz).

We modified the original structure of the pipeline provided by IntroVerse and included the following data filters. Samples from fresh frozen preserved tissues were prioritised. On this basis, samples from *“Brain - Cortex”* and *“Brain - Cerebellum”* tissues were discarded as well as all sex-specific tissues and tissues with less than 70 samples (*e.g. Bladder, Cells - Leukaemia cell line (CML), Cervix - Ectocervix, Cervix - Endocervix, Fallopian Tube and Kidney - Medulla*). In addition, only samples presenting an RNA Integrity Number (RIN) higher or equal to 6 were included in this study, as any more stringent RIN thresholds would have reduced excessively the number of samples available for study:

*RIN* ≥8, *NSamplesAvailable* = 4, 170;

*RIN* ≥7, *NSamplesAvailable* = 9, 494;

*RIN* ≥6, *NSamplesAvailable* = 14, 408.

In addition, we discarded n=555 annotated introns reported to be spliced by the minor spliceosome^88^ and n=9,252 novel donor and novel acceptor junctions linked to them. We discarded these minor introns because, even though they represent less than 1% of all intervening sequences in the human genome, their consensus splicing sequences differ considerably from the consensus sequences of the human introns targeted by the major spliceosome^89^. To avoid biases derived from these differences, introns targeted by the minor spliceosome were discarded.

The abovementioned modifications resulted in a new relational database, namely *Splicing* intron database, which included a set of 324,956 annotated introns (Ensembl-v105) and a linked set of 3,865,268 novel junctions, covering 14,408 different human samples and 42 human tissues (**Extended Data Fig. 1a,b**). All types of exon-exon junction reads were considered (jxn_format = c(“ALL”), recount3::create_rse_manual() function, Bioconductor R package recount3 version 1.0.7, https://bioconductor.org/packages/release/bioc/html/recount3.html).

### Calculating the contamination rates across multiple version of the Ensembl transcriptome

Split reads were first annotated based on the reference transcriptome Ensembl-v97 (v97) released in July 2019 and using the Bioconductor R package *dasper* version 1.4.3 (https://bioconductor.org/packages/release/bioc/html/dasper.html).

Per each tissue, we compared the introns that had been classified as novel donor or novel acceptor junctions using v97 but were also re-annotated as annotated introns in the Ensembl-v105 (v105), and used them as a measure of *“contamination”*. To create a normalised measure of contamination rates across the tissues, we divided the number of novel junctions in v97 that had been classified as annotated introns in v105 by the total number of novel junctions that had maintained annotation category between the two aforementioned Ensembl versions. Finally, we converted the resulting ratio figure into a percent.

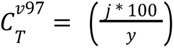

Let *j* denote the total number of unique novel donor and novel acceptor junctions in v97 that had been re-classified as annotated introns in v105. Let *y* denote the total number of unique novel donor and novel acceptor junctions in v97 that had maintained annotated category in v105. Let *T* denote the tissue studied.

This approach was mirrored to re-annotate all split reads from the frontal cortex brain tissue using four different Ensembl versions v76, v81, v90 and v104 published in July 2014, July 2015, July 2017 and March 2021, respectively. Contamination rates in each Ensembl version were again calculated using v105 as the reference annotation.

### Calculating the percentage of unique novel junctions and novel read counts

Focusing on the novel donor category, the percentage of unique novel donor junctions in a given tissue was calculated by dividing the cumulative number of unique novel donor junctions across all samples of the studied tissue by the total number of unique annotated introns, novel donor and acceptor junctions found across the same set of samples. Finally, we converted the resulting ratio to a percentage.

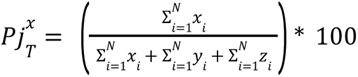

Let *x* denote the total number of unique novel donor junctions within one sample of the tissue *T* studied. Let *y* denote the total number of unique novel acceptor junctions within one sample of tissue *T*. Let *z* denote the total number of unique annotated introns within one sample of tissue *T*. Let *N* denote the total number of samples studied of tissue *T*. Let *T* denote the tissue studied.

We mirrored the method detailed above to calculate the percentage of unique annotated introns and the percentage of unique novel acceptor junctions within a tissue.

Similarly, focusing on the novel donor category, the percentage of novel donor read counts in a given tissue was calculated by dividing the cumulative number of novel donor reads counts by the total number of reads mapping to annotated introns, novel donor and acceptor junctions across all samples of the tissue studied. The resulting ratio was multiplied by 100 to create a percentage.

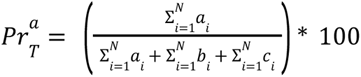

Let *a* denote the total number of read counts that all novel donor junctions presented within one sample of tissue *T*. Let *b* denote the total number of read counts that all novel acceptor junctions presented within one sample of tissue *T*. Let *c* denote the total number of read counts that all annotated introns presented within one sample of tissue *T*. Let *N* denote the total number of samples studied of tissue *T*. Let *T* denote the tissue studied.

We mirrored the formula above to calculate the percentage of annotated introns and novel acceptor read counts within a tissue.

### MaxEntScan score analyses

The MaxEntScan^50^ (MES) algorithm (version 1.0, RRID:SCR_016707, http://genes.mit.edu/burgelab/maxent/Xmaxentscan_scoreseq.html) was applied to score the 9 bp sequence at the 5’ss and the 23 bp sequence at the 3’ss of each annotated intron and novel junction stored on the *Splicing* intron database. The higher is the MES score assigned to a given sequence, the stronger that sequence will be considered, as it will be more closely related to a real annotated splice site. To investigate the differences in the strength implied by each novel splice site and the analogous annotated splice site of its paired annotated intron, we obtained the delta values of their MES scores.

Focusing on the novel donor junctions, the delta MES (Δ*MES*5*ss*) was calculated by obtaining the difference between the MES score assigned to the 9 bp sequence at the 5’ss of the annotated intron (*MESann*5*ss*) minus the MES score assigned to the 9 bp sequence at its paired 5’ss of the novel donor junction (*MESdon*5*ss*):

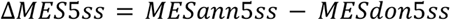

Similarly, to calculate the delta MES at the acceptor sites (Δ*MES*3*ss*), we obtained the difference between the MES score assigned to the 23 bp sequence at the 3’ss of the annotated intron (*MESann*3*ss*) minus the MES score assigned to the 23 bp sequence at the 3’ss of its linked novel acceptor junction (*MESacc*3*ss*):

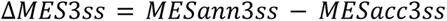

### Calculating the Distances and Modulo3

Per each tissue analysed, we calculated the distances lying between each novel splice site and the analogous annotated splice site of their linked annotated intron. Focusing on the novel donor junctions, we obtained the distances in bp lying between the novel 5’ss of each novel donor junction and the annotated 5’ss of their linked annotated intron. We repeated this process to calculate the distances at 3’ss.

Distances in bp were calculated by following a 0-based genomic-interval approach, as we required splicing to occur at precise annotated genomic coordinates to consider splicing as accurate. For instance, focusing in a novel donor junction whose novel 5’ss is located at the *gcNovel* genomic coordinate, the distance lying between *gcNovel* and the 5’ss of its linked annotated intron (*gcIntron*) can be expressed as:

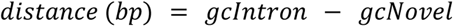

Let *gcIntron* denote the genomic coordinate corresponding to the 5’ss of the annotated intron *Intron* (Ensembl-v105). Let *gcNovel* denote the genomic coordinate corresponding to the 5’ss of the novel donor *Novel* attached to the annotated intron *Intron*. Let *distance* denote the difference in bp between the two genomic positions *gcIntron* and *gcNovel* within the same strand.

The formula above was mirrored to calculate the distances lying between each novel acceptor junction and its linked annotated intron.

For the Modulo3 analysis, we restricted the analysis to the annotated introns belonging to transcripts categorised as MANE Select^90^, as these represented exact matches in exonic regions between Refseq transcript and the Ensembl/GENCODE. Only the novel junctions located less than 100 bp apart from annotated splice sites were considered. This filter increased the confidence for the novel products to be located within the adjacent exon and intron sequences, as the average exon size corresponds to 120 bp^91^, whereas the mode, median and average length of the annotated introns corresponded to 88 bp, 1,945 bp and 8,388 bp, respectively (**Extended Data Fig. 10**).

### Calculating the Mis-Splicing Ratio measures

Focusing on the mis-splicing activity at the 5’ss of a given annotated intron, the *MSR_D_* measure represented the ratio of cumulative number of novel donor reads found across all annotated reads linked to the annotated intron of interest and across all samples of a given tissue.

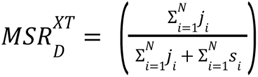

Let *j* denote the total number of novel donor junction reads assigned to the annotated intron *X* within one sample of the tissue *T*. Let *s* denote the total number of annotated intron reads for the same intron, *X*, within the same sample of study. Let *N* denote the total number of samples studied from the tissue *T*.

The *MSR_D_* and *MSR_A_* measures were normalised to values between 0 and 1. Hence, *MSR_D_* = 0 represents accurate splicing at the 5’ss of a given annotated intron, whereas *MSR_D_* = 1 represents complete mis-splicing at the 5’ss of a given annotated intron.

### Calculating the Transcript per Million (TPM) measure

To calculate the Transcript Per Million value corresponding to a particular gene within a set of samples, we obtained the total number of read counts that each gene presented per sample and divided this number by the length of the gene in kilobases, namely RPK number. We then summed all the RPK values calculated per sample and divided this number by 1,000,000. This division created a figure named the per million scaling factor. Finally, the RPK values were divided by the “per million” scaling factor.

### Using linear models to predict the *MSRD* and the *MSRA* measures

For each GTEx tissue, we built two linear regression models to discover the characteristics influencing the rate of mis-splicing at the 5’ss (*MSR_D_*) and 3’ss (*MSR_A_*) of the set of annotated introns studied. To reduce any biases regarding the number of introns considered per tissue, we only included the annotated introns that were commonly found across all tissues. These were n=151,729 common annotated introns.

As predictors, we included covariates encompassing diverse gene and intron-level features that were provided by the IntroVerse database. The gene-level covariates included (1) the gene length in bp, (2) the median transcript per million (TPM) of the gene calculated across the samples of each tissue studied **(Methods)**, (3) the total number of transcripts of the gene (Ensembl-v105) and (4) the percent of protein-coding transcripts in which the assessed intron may appear. The intron-level covariates included (1) the MES^50^ scores of the sequences overlapping the 5’ss and 3’ss, (2) the intron length in bp, (3) the mean interspecies conservation score across 20 species^92^ (phastCons20) overlapping the proximal intronic sequences, and (4) the mean context-dependent tolerance score (CDTS) scores^66^ overlapping the aforementioned proximal intronic sequences. The proximal intronic sequences were defined as the −5/+35 bp sequence window around the exon-intron junction at the 5’ss of each annotated intron, and the −35/+5 bp sequence neighbouring the intron-exon junction at the 3’ss of each annotated intron. The mean phastCons20 scores represented the probability of negative selection based on the number of substitutions occurring across 20 species (human, 16 primates, dog, mouse and tree shrew) during evolution. The CDTS scores represented the sequence constraint across humans.

Focusing on the prediction of the *MSR_D_*, the formula used to build the corresponding linear model is shown below:

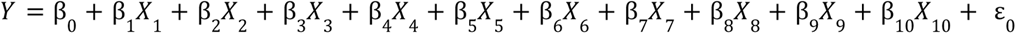

where the dependent variable corresponded to: *Y* = *MSR_D_* and the independent variables were:

*X*_1_ = *gene length*;

*X*_2_ = *median TPM of the gene across the cluster of samples studied*;

*X*_3_ = *number of transcripts in annotation of the gene*;

*X*_4_ = *percentage of protein coding transcripts in which the intron may appear*;

*X*_5_ = *MES of the* 5’*ss of the intron*;

*X*_6_ = *MES of the* 3’*ss of the intron*;

*X*_7_ = *Mean PhastCons*20 *score of the sequences neighbouring the* 5’*ss of the intron*;

*X*_8_ = *Mean PhastCons*20 *score of the sequences neighbouring the* 3’*ss of the intron*;

*X*_9_ = *Mean CDTS score of the sequences neighbouring the* 5’*ss of the intron*;

*X*_10_ = *Mean CDTS score of the sequences neighbouring the* 3’*ss of the intron*;

*ε*_0_ = *N*(0, *sigma*^2^)

We mirrored the formula above to predict the *MSR_A_* value of the introns studied.

In total, 84 linear regression models were generated. We created 1 linear model to predict the *MSR_D_* value of the set of n=151,729 annotated introns studied and another linear model to predict the *MSR_A_* value of the same set of introns in each of the 42 tissues considered. P-values were FDR-adjusted, producing *q* values. β values were then grouped by MSR, generating two independent distributions that represented the tissue variability in the effect that each covariate produced in the prediction of the mis-splicing rates at the 5’ss (*MSR_D_*) and 3’ss (*MSR_A_*) of the set of common n=151,729 introns studied across the tissues.

### Assessing the levels of mis-splicing in sun-exposed versus non-sun-exposed skin tissues

Using data from the *Splicing* database, we selected all annotated introns from the *“Skin - Sun Exposed (Lower leg)”* and the *“Skin - Not Sun Exposed (Suprapubic)”* body sites, and evaluated their differences in mis-splicing rates at their 5’ss (*MSR_D_*) and 3’ss (*MSR_A_*).

*“Skin - Not Sun Exposed (Suprapubic)”* originally presented a total of n=250,948 annotated introns, whereas *“Skin - Sun Exposed (Lower leg)”* presented n=251,769 annotated introns. We next obtained the common annotated introns overlapping both tissues. This reduced both distributions to a set of n=232,885 unique common annotated introns. In addition, to reduce any potential biases derived from differences in the sequencing depth levels of the two sets of samples, we subsampled the set of n=232,885 annotated introns between the two skin body sites by pairing them by mean coverage similarity. The intron pairing was performed by restricting the maximum difference in log10 mean coverage to 0.005 reads (we used the matchit() function, MatchIt R package^93^, version 4.4.0, https://cran.r-project.org/web/packages/MatchIt/vignettes/MatchIt.html). This pairing reduced both distributions of annotated introns to a total of n=227,038.

Finally, we obtained the *MSR_D_* and *MSR_A_* values from each of the n=227,038 annotated introns of either *“Skin - Sun Exposed (Lower leg)”* and *“Skin - Not Sun Exposed (Suprapubic)”*. To test for any significant differences in the median distribution of these two mis-splicing ratio measures between the two skin body sites, we used a paired Wilcoxon signed rank test function with continuity correction (wilcox_test() function, R package rstatix^94^ version 0.7.1, RRID:SCR_021240, https://CRAN.R-project.org/package=rstatix).

### Analysing shRNA knockdown of RBPs followed by RNA-sequencing data from ENCODE

From the list of 356 RBPs published by Nostrand et al. in 2020^54^, we selected 115 RBPs that had been functionally categorized as splicing regulation, spliceosome or exon-junction complex by the authors. We also downloaded a second list of 118 human genes published by the Reactome project that had been classified as involved in NMD processes [R-HSA-927802, NMDv3.7, Browser v82]. In total, 233 genes were considered for study.

From the 233 genes initially considered, only 56 had shRNA knockdown followed by RNA-sequencing data available on the ENCODE platform. For each of these 56 genes, the same number of experiments was used. Experiments were chosen based on similarity of metadata and design. Briefly from ENCODE: (1) 4 experiments with RNA-seq data available on K562 and HepG2 cells treated with an shRNA knockdown against a given gene, and (2) 4 control shRNA experiments against no target gene were chosen for each gene. Details and metadata of all the shRNA knockdown experiments downloaded from ENCODE are shown in **(Supplementary Table 12)**.

A total of 8 alignment BAM files (GRCh38 v29) were downloaded per gene, each one corresponding to a different ENCODE experiment. In total, 448 BAM files were downloaded from the ENCODE platform. To extract the splicing junctions from the BAM files to a BED12 format, we made use of the command *‘regtools junction extract’* made available through the regtools software package (version 0.5.2, http://regtools.org/). We required a minimum and maximum intron size of 25 and 1,000,000 bp respectively, and the strand information was provided by the aligner. Prior to the extraction, alignment reads were sorted and indexed using the commands *‘samtools sort’* and *’samtools index’*, both made available through the SAMTOOLS^95^ software (version 1.16.1, RRID:SCR_002105, http://htslib.org/).

We then applied a similar data analysis to the one originally published by IntroVerse (**Methods**), and created a separate database for each ENCODE shRNA knockdown project, in which samples were clustered following a case/control grouping criteria. Case samples corresponded to the experiments in which a gene had been targeted for knockdown, whereas control samples corresponded to untreated controls in which no gene had been targeted. To account for any differences in read-depth or RIN numbers across the different samples and experiments compared, we only considered the annotated introns that were common across all samples and projects. These represented a total of n=109,950 annotated introns (Ensembl-v105). For more details about how the MES, distances, Modulo3 and MSR measures were calculated, please refer to the following sections (***Methods: “MaxEntScan score analyses”, “Calculating the Distances and Modulo3”, and “Calculating the Mis-Splicing Ratio measures”***).

To detect any significant differences for each gene between case versus control samples in the MSR median values of the common introns, we made use of the wilcox_test function (R package rstatix^94^ version 0.7.1, RRID:SCR_021240, https://CRAN.R-project.org/package=rstatix). Focusing on the study of mis-splicing at the 5’ss, the null hypothesis (*H*_0_) tested corresponded to: *“the set of common introns studied do not present different MSR_D_ median values in case versus control samples”*. On the contrary, the alternative hypothesis (*H*_1_) tested corresponded to: *“the set of common introns studied present higher MSR_D_ median values in case versus control samples”.* We then repeated these hypothesis to test for differences in the *MSR_A_* values. A total of 112 Wilcoxon tests were run, one per ENCODE knockdown project and splice site. The p-values obtained from each test were adjusted using the Bonferroni correction method. In those cases in which the alternative hypothesis (*H*_1_) was accepted, we calculated the probability of superior MSR outcome in case vs control samples by using the function wilcox_effsize() (R package rstatix, version 0.7.1, RRID:SCR_021240, https://CRAN.R-project.org/package=rstatix).

### Knockdown efficiency extraction

To obtain a measurement of the knockdown efficiency for each ENCODE experiment, we identified a *’biosample preparation and characterization’* document attached to 46 out of the 56 studied genes. The efficiency is calculated by comparing protein levels in control and knockdown cells using a western blot analysis, and reported in figures embedded in the document. To extract the figures, we made use of the *’fitz’* module available from the python package PyMuPDF^96^ (version 1.21.1, https://github.com/pymupdf/PyMuPDF). We employed the Tesseract-OCR (Optical Character Recognition) algorithm, available through the python package pytesseract (version 0.3.10, https://pypi.org/project/pytesseract/) to extract the text from the images. To ensure high accuracy in the image to text conversion, figures were: (1) cropped to only contain the depletion percentages, and (2) resized to a lower resolution to better match the training data of the OCR algorithm. No additional configuration was specified to the Tesseract-OCR engine. A perfect accuracy was observed when tested in 15% of the samples, and outliers were manually verified. The final reported knockdown efficiency is the average of the measurements for all four samples.

### RBP expression levels across tissues

We visualised the gene expression for 115 important spliceosomal RBP genes across all 42 GTEx v8 tissues, deriving the RBP gene list from Van Nostrand et al. 2020. In order to gauge cross-tissue variation in expression for each gene, the following calculations were performed on a per-gene basis. Firstly, we obtained the cross-sample expression for each tissue, identifying the tissue with the median expression value, namely tissue Y. Next, we calculated the log2 fold-change in expression for each of the remaining 41 tissues in relation to expression in tissue Y. Finally, the log2 fold-change expression values for each gene were visualised as a heatmap facetted by gene functional group. The functional categories used were “Splicing Regulation”, “Spliceosomal” and “Exon-Junction complex”, obtained from Van Nostrand et al. 2020. The code to reproduce this analysis can be accessed at https://github.com/ainefairbrother/RBP_expression_analysis (version 1.0.0, DOI: 10.5281/zenodo.7736907).

### Changes in RBP expression levels with age in brain tissue

We downloaded the read counts from all genes expressed within the Brain GTEx v8 tissue from the recount3 project and using the function *create_rse_manual()* (R package recount3, version 1.0.7, https://bioconductor.org/packages/release/bioc/html/recount3.html). Raw counts were transformed using the function *transform_counts()* (R package recount3, version 1.0.7, https://bioconductor.org/packages/release/bioc/html/recount3.html). To calculate the gene expression within each sample, we normalized the read counts using the TPM method **(Methods: Calculating the Transcript per Million (TPM) measure)**. TPM data was used in this analysis because all samples had been obtained from the same tissue (i.e. brain), and all samples had been sequenced using the same library protocol, polyA-selection, reducing the risk of misleading TPM comparisons^97^.

To know whether the expression levels of the 116 RBPs involved in splicing regulation, spliceosomal and exon-junction recognition^54^ functions were affected by age in brain tissue samples, we built a linear regression model per RBP. The independent variable to predict corresponded to the TPM value of each RBP in each sample. The dependent variables corresponded to a set of covariates providing information about the sample: age, center, gebtch, gebtchd, nabtc, nabtchd, nabtcht, hhrdy, sex and rin. These covariates were chosen by following the results from the principal component analysis (PCA) published by Fairbrother-Browne, A. et. al^98^ using data from GTEx v6. Some of these covariates were categorical, so we transformed them into numerical values prior inclusion to the linear models.

In total, 116 linear models were run, one per RBP studied. Each linear model was built to predict 2,363 TPM values per RBP, equating to the total number of brain samples available. P-values produced by each linear model were corrected for multiple testing using the Benjamini-Hochberg method, producing *q* values. Finally, in those cases in which the age covariate produced a negative estimate value in the prediction of the TPM for a given RBP, it was considered that age negatively affected the expression levels of that given RBP across the set of brain samples studied.

### Age stratification and sample clustering

GTEx tissues were grouped by tissue following the original classification made by recount3^83^ (**Supplementary Table 13**). Samples from each body region were then binned by age within one of these three categories *“20-39”, “40-59”* and *“60-79”*. Only the body sites presenting a minimum of 75 samples, equating to at least 25 samples per age category, were considered. These were 18 body sites in total: “ADIPOSE TISSUE”, “ADRENAL GLAND”, “BLOOD”, “BLOOD VESSEL”, “BRAIN”, “COLON”, “ESOPHAGUS”, “HEART”, “LUNG”, “MUSCLE”, “NERVE”, “PANCREAS”, “SALIVARY GLAND”, “SKIN”, “SMALL INTESTINE”, “SPLEEN”, “STOMACH” and “THYROID”.

To account for differences in the RIN numbers presented by the samples grouped in each age category, we down sampled the clusters *“40-59”* and *“60-79”* to meet similarity with the *“20-39”* group, as the overall sample size of the latter was always lower than the two former categories across all body sites studied. The sample pairing was performed only when two samples from each age group presented a maximum difference of 0.05 in their RIN numbers (matchit(), MatchIt R package^93^, version 4.4.0, https://cran.r-project.org/web/packages/MatchIt/vignettes/MatchIt.html) (**Extended Data Fig. 7, Supplementary Table 14**).

We then applied our modified version of the pipeline published by IntroVerse ***(Methods: RNA-sequencing data download and processing)*** and created a relational database to study the changes occurring in the splicing activity of the n=308,717 annotated introns (Ensembl-v105) that were found across the three age categories and 18 body sites studied. We named it the *“Age-Stratification”* intron database (**Extended Data Fig. 8**), and it contained a total of 308,717 annotated introns, from which 228,534 presented evidence of at least one type of mis-splicing event. It also included 719,069 novel donor and 999,041 novel acceptor junctions, covering 199,191 transcripts, 30,580 genes and 6,111 samples from 40 body sites and 18 tissues.

To study the effect size of mis-splicing produced by age at the 5’ss and 3’ss of the annotated introns stored on the *“Age Stratification”* intron database, we made use of their *MSR_D_* and *MSR_A_* values. To reduce any biases in the number of annotated introns considered in this comparative analysis across multiple body sites, we only included the introns that were common across the three age categories and all 18 tissues studied. These were a total of n=137,713 annotated introns. Then, to further reduce the likelihood of including borderline samples between the three age groups, we only considered samples from the two most extreme age clusters “20-39” and “60-79”.

Focusing on the 5’ss of the n=137,713 common annotated introns overlapping the “20-39” and “60-79” age groups, we calculated the Wilcoxon effect size that the covariate *age* (i.e. “20-39” and “60-79”) was producing over their *MSR_D_* values. We then repeated this approach to measure the effect size between age and the mis-splicing activity at the 3’ss (*MSR_A_*) of the same set of n=137,713 annotated introns. In both cases, we made use of the function wilcox_effsize (R package rstatix^94^, version 0.7.0, RRID:SCR_021240, https://CRAN.R-project.org/package=rstatix).

### Gene Ontology enrichment of genes containing introns with increasing levels of MSR values in ageing samples

Using data from the *‘Age-Stratificatioń* database, we selected all introns overlapping the three age categories for the brain tissue. These were n=206,067 annotated introns. To assess any changes occurring in their splicing activity, we compared their *MSR_D_* and *MSR_A_* measures and evaluated the changes occurring as the age of each cluster increased. Focusing on the *MSR_D_* value, we selected the introns presenting increasing levels of mis-splicing with age at their 5’ss (*MSR_D_* 20-39yrs < 40-59yrs < 60-79 yrs). We mirrored this approach focusing on their *MSR_A_*.

Then, we obtained the gene symbol of all introns showing increasing *MSR_D_* and/or *MSR_A_* values with age. These were a total of n=8,117 unique genes. Using as background the list of all genes (n=19,140) parenting the complete set of annotated introns found across brain sites, we ran a GO and KEGG enrichment analysis of the set of n=8,117 unique genes. For the GO enrichment analysis, we used the R function enrichGO (R package clusterProfiler, version 3.18.1, RRID:SCR_016884, http://yulab-smu.top/biomedical-knowledge-mining-book/clusterprofiler-go.html). For the KEGG enrichment analysis, we used the R function enrichKEGG (R package clusterProfiler, version 3.18.1, RRID:SCR_016884, http://yulab-smu.top/biomedical-knowledge-mining-book/clusterprofiler-kegg.html?q=enrichKEGG#clusterprofiler-kegg-pathway-ora).

### RBP cell-type enrichment calculation

We used Expression Weighted Cell Type Enrichment (EWCE)^99^ (https://bioconductor.org/packages/EWCE) to determine whether genes involved in splicing regulation have higher expression within particular brain-related cell types than would be expected by chance. We used two gene lists: (i) a list of 115 RBPs that had been functionally categorized as splicing regulation, spliceosome or exon-junction complex by Nostrand et al.^54^, and (ii) a list of 118 human genes published by the Reactome project that had been classified as involved in NMD processes [R-HSA-927802, NMDv3.7, Browser v82]. In total, 233 genes were considered for study.

Our aim was to evaluate the average level of expression of the 233 aforementioned genes within the data set Human Multiple Cortical Areas SMART-seq, which includes single-nucleus transcriptomes from 49,495 nuclei across multiple human cortical areas. These data are freely available through the Allen Brain Atlas^62^ data portal (https://portal.brain-map.org/atlases-and-data/rnaseq). To achieve this aim, we first downloaded the *EWCE* docker image (https://hub.docker.com/r/neurogenomicslab/ewce), which includes the EWCE^100^ R package (version 0.99.3, https://bioconductor.org/packages/release/bioc/html/EWCE.html). Second, we downloaded the single-nucleus transcriptomes from 49,495 nuclei across multiple human cortical areas from https://portal.brain-map.org/atlases-and-data/rnaseq/human-multiple-cortical-areas-smart-seq. We made use of the matrices including exon and intron counts. For this analysis, all brain regions sampled were included, which corresponded to:

- Middle temporal gyrus (MTG)
- Anterior cingulate cortex (ACC; also known as the ventral division of medial prefrontal cortex, A24)
- Primary visual cortex (V1C)
- Primary motor cortex (M1C) - upper (ul) and lower (lm) limb regions
- Primary somatosensory cortex (S1C) - upper (ul) and lower (lm) regions
- Primary auditory cortex (A1C)

Then, we generated the cell type annotations:

- Level 1: Allen Brain Atlas provided a class and subclass label. Class had only 3 levels (GABAergic, glutamatergic and non-neuronal), thus instead we used the subclass label, which subdivided glutamatergic neurons into 7 subtypes, GABAergic neurons into 5 subtypes, and non-neuronal cell types into Astrocyte, Endothelial, Microglia, Oligodendrocyte, OPC, Pericyte, VLMC. As the number of endothelial cells (n = 70), pericytes (n = 32) and VLMC (n = 11) nuclei was low, these were merged into the class “vascular cell”.
- Level 2: used the original clusters defined by the Allen Brain Atlas.

A total of 1,985 nuclei were labelled as “outlier calls” and were removed during generation of the celltype dataset. We used the function *fix_bad_hgnc_symbols()* (R package EWCE, version 0.99.3, https://bioconductor.org/packages/release/bioc/html/EWCE.html) to remove any symbols from the gene-cell matrix that were not official HGNC symbols. A total of 30,792 genes were retained.

We then used the function *drop_uninformative_genes()* (R package EWCE, version 0.99.3, https://bioconductor.org/packages/release/bioc/html/EWCE.html), which removes *“uninformatic genes”* to reduce compute time in subsequent steps. The following steps were performed:

- Drop non-expressed genes (n=1,263). This step removed the genes that are not expressed across any cell types.
- Drop non-differentially expressed genes (n=6,304), which removes genes that are not significantly differentially expressed across level 2 cell types with an adjusted p-value threshold of 1e-05.

Finally, we used the function *generate_celltype_data()* from the R package EWCE (version 0.99.3, https://bioconductor.org/packages/release/bioc/html/EWCE.html) to generate the celltype dataset. This dataset can be accessed at: https://github.com/RHReynolds/MarkerGenes (version 0.99.1, DOI: 10.5281/zenodo.6418604).

In a separate analysis run in R 4.2.0, we used this cell type data reference in EWCE. The goal of this analysis was to determine whether the genes of interest had significantly higher expression in certain cell types than might be expected by chance. Bootstrap gene lists controlled for transcript length and GC-content were generated with EWCE iteratively (n=10,000) using *“bootstrap_enrichment_test()”* function. In brief, this function takes the inquiry gene list and a single cell type transcriptome data set and determines the probability of enrichment of this list in a given cell type when compared to the gene expression of bootstrapped gene lists; the probability of enrichment and fold-change of enrichment are the returned. P-values were corrected for multiple testing using the Benjamini-Hochberg method. The code, plotting and library versions used for this analysis can be accessed at: https://github.com/mgrantpeters/RBP_EWCE_analysis (version 1.0, DOI: 10.5281/zenodo.7734035).

### Code availability

The repository https://github.com/SoniaRuiz/splicing-accuracy-manuscript (version 1.0.1, DOI: 10.5281/zenodo.7717150), contains the code to (1) generate the two *“Splicing”* and *“Age Stratification”* intron databases, and (2) replicate all the main and supplementary analyses and figures described in this manuscript.

The code to calculate the expression levels of the RBPs known to contribute to splicing and its regulation across body sites can be accessed at https://github.com/ainefairbrother/RBP_expression_analysis (version 1.0.0, DOI: 10.5281/zenodo.7736907).

The code to obtain the metadata from an ENCODE experiment can be accessed at https://github.com/guillermo1996/ENCODE_Metadata_Extraction (version 1.0.2, DOI: 10.5281/zenodo.7733986).

The code to reproduce the shRNA knockdown comparison between cases and control ENCODE experiments is available at: https://github.com/guillermo1996/ENCODE_Splicing_Analysis (version 1.0.1, DOI: 10.5281/zenodo.7733984).

The code to reproduce the cell type specificity analysis of the set of RBPs known to contribute to splicing and its regulation, and using as reference the drop-seq data from multiple cortical regions (Allen Brain Atlas) is available at: https://github.com/mgrantpeters/RBP_EWCE_analysis (version 1.0, DOI: 10.5281/zenodo.7734035).

The code to generate the celltype dataset using the function generate_celltype_data() from the R package EWCE, can be accessed at: https://github.com/RHReynolds/MarkerGenes (version 0.99.1, DOI: 10.5281/zenodo.6418604).

The Supplementary Tables are available at https://zenodo.org/record/7732872 (DOI: 10.5281/zenodo.7732872).

## Extended Data Figures

**Extended Data Fig. 1:**
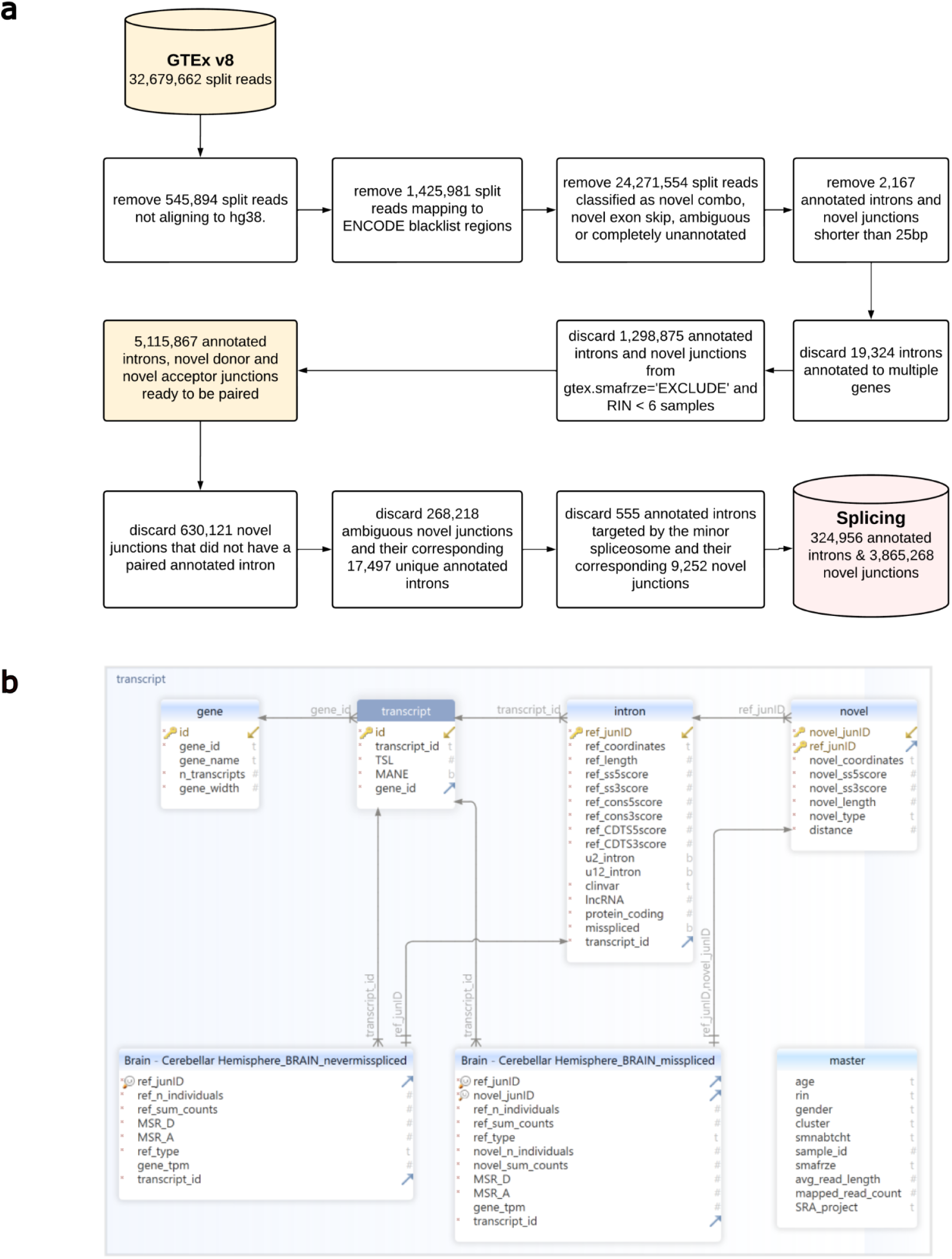
Generation of the Splicing database. **a.** Overview of the quality-control steps applied to the dataset of exon-exon junctions provided by IntroVerse to produce the Splicing database. **b.** SQL schema of the Splicing database.

**Extended Data Fig. 2:**
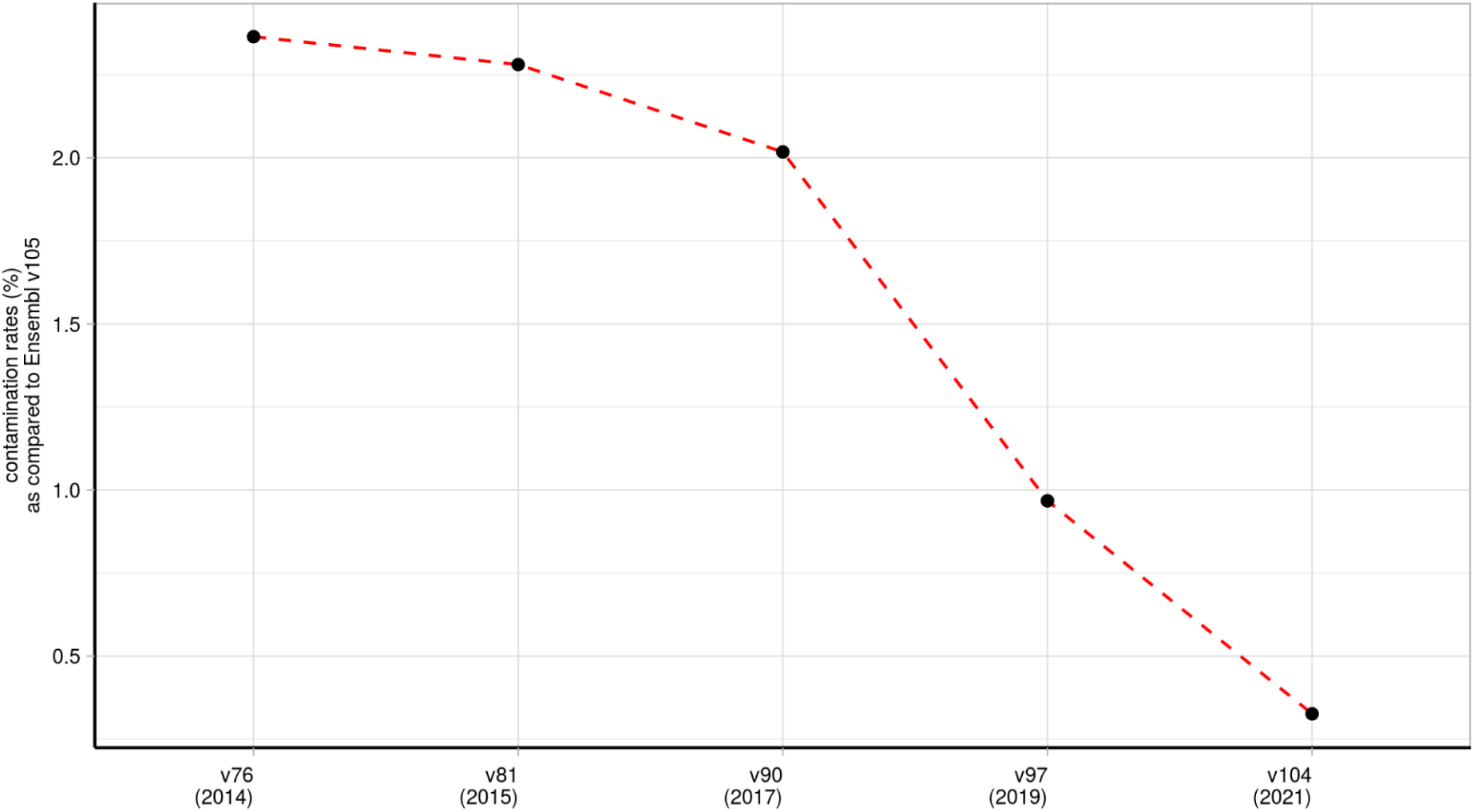
Contamination rates in Ensembl v97 as compared to Ensembl v105 in samples from the frontal cortex brain tissue. Each point represents the percentage of novel junctions in each Ensembl version (x-axis) that entered annotation as annotated introns in Ensembl v105 (2021).

**Extended Data Fig 3:**
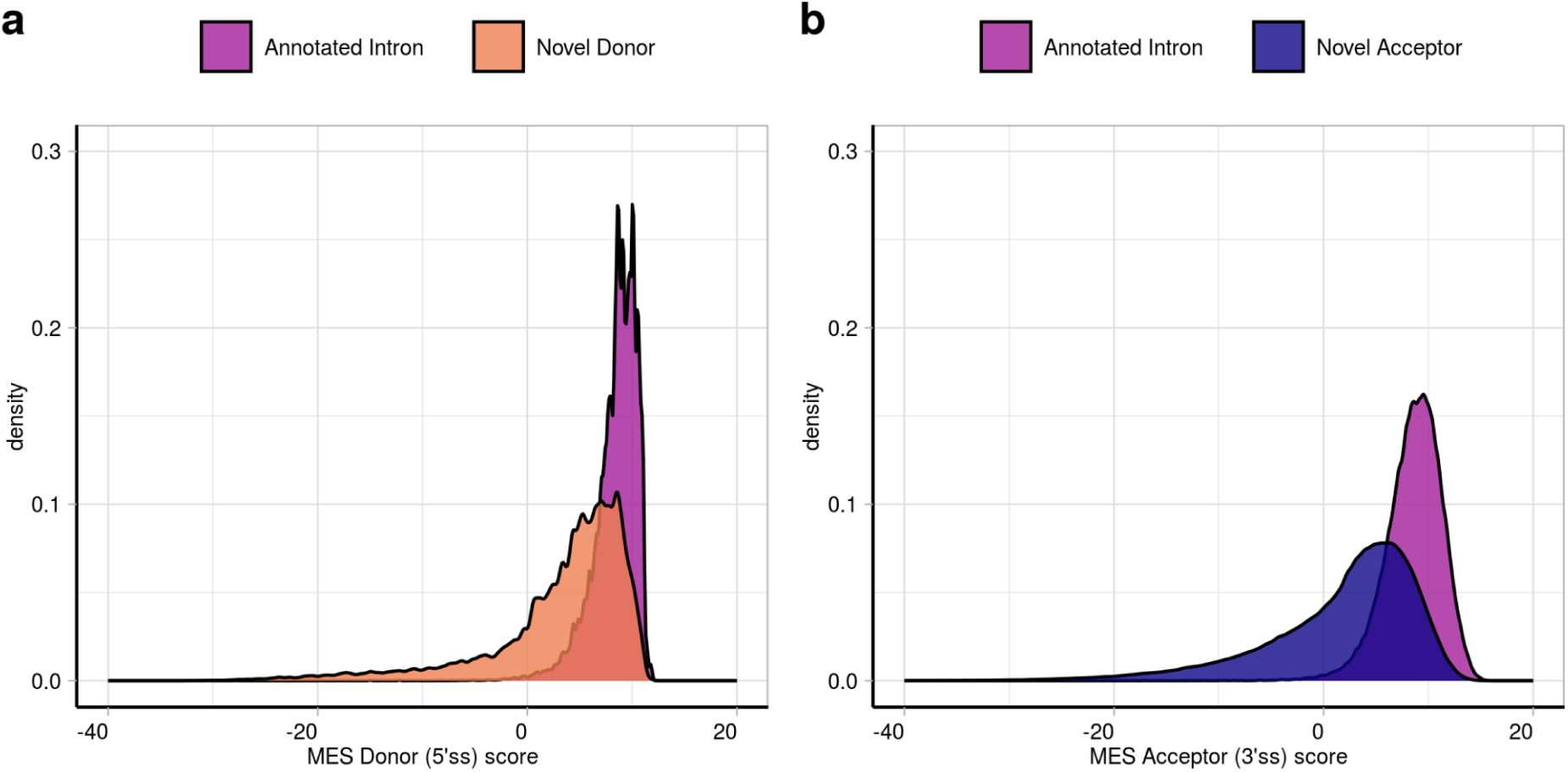
Comparison of the MES scores assigned to the donor and acceptor splice sites of the annotated introns compared to the MES scores assigned to donor and acceptor splice sites of their paired novel junctions. **a.** MaxEntScan scores assigned to the donor splice site (i.e. 5’ss) of all novel donor junctions found across all tissues (in orange) and compared with the MES scores assigned to the annotated donor splice site of their linked annotated introns (in dark pink). **b.** MaxEntScan scores assigned to the acceptor splice site (i.e. 3’ss) of all novel acceptor junctions found across all tissues (in dark blue) and compared with the MES scores assigned to the annotated acceptor splice site of their linked annotated introns (in dark pink).

**Extended Data Fig 4:**
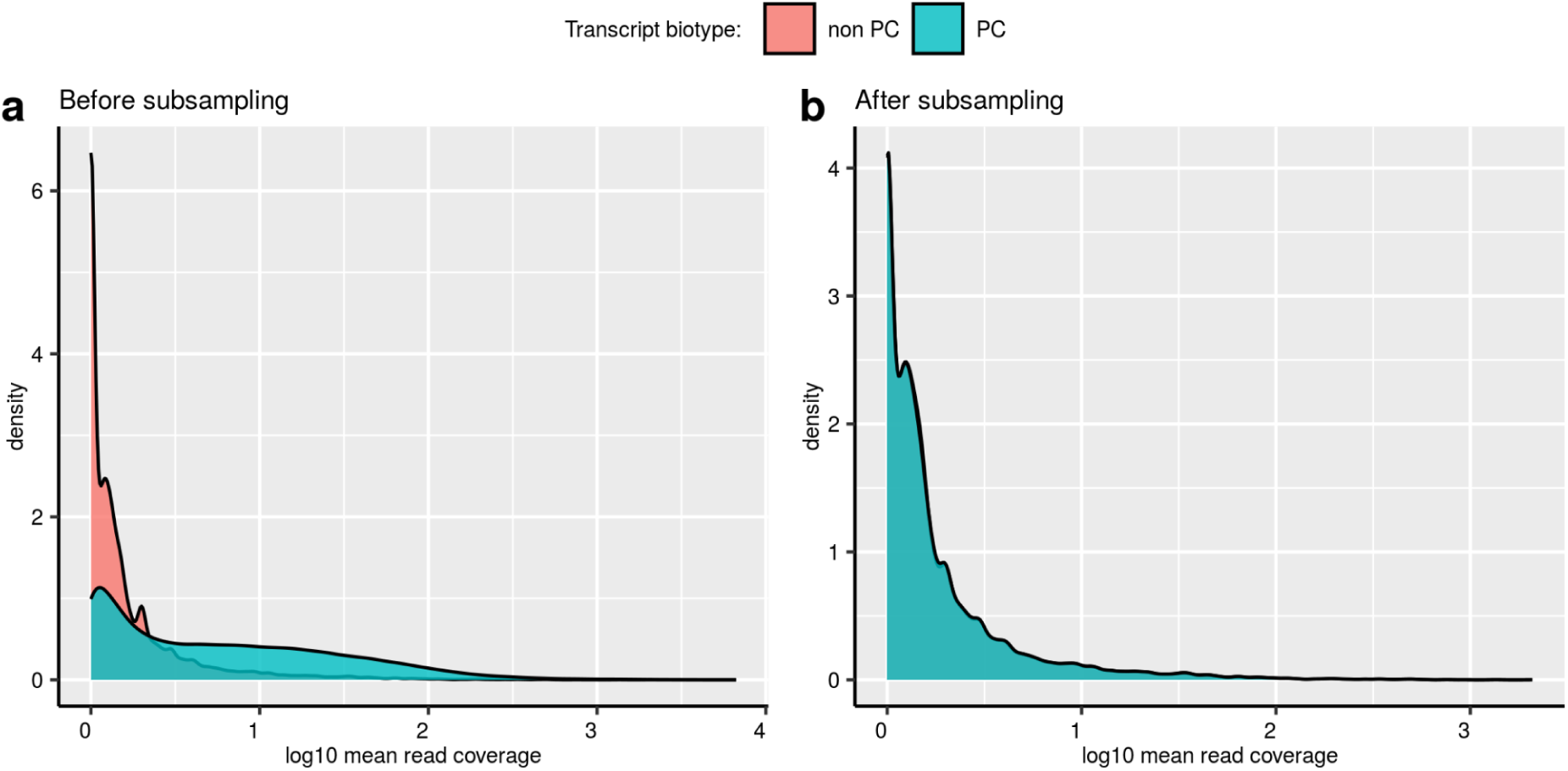
Mean read coverage of the annotated introns found across the samples of the frontal cortex brain tissue. **a.** Mean read coverage of the annotated introns from non-protein-coding transcripts compared to annotated introns from protein-coding transcripts (PC) before subsampling by mean read coverage similarity. Data from frontal cortex brain tissue. Only annotated introns presenting a maximum difference of 0.005 in their mean read coverage between the two data sets were considered, paired and kept for downstream analyses. **b.** Mean read coverage of the annotated introns from non-protein-coding transcripts (non PC) compared to protein-coding transcripts (PC) after subsampling by mean read coverage similarity. Data from frontal cortex brain tissue. Only annotated introns presenting a maximum difference of 0.005 in their mean read coverage between the two data sets were considered, paired and kept for downstream analyses.

**Extended Data Fig. 5:**
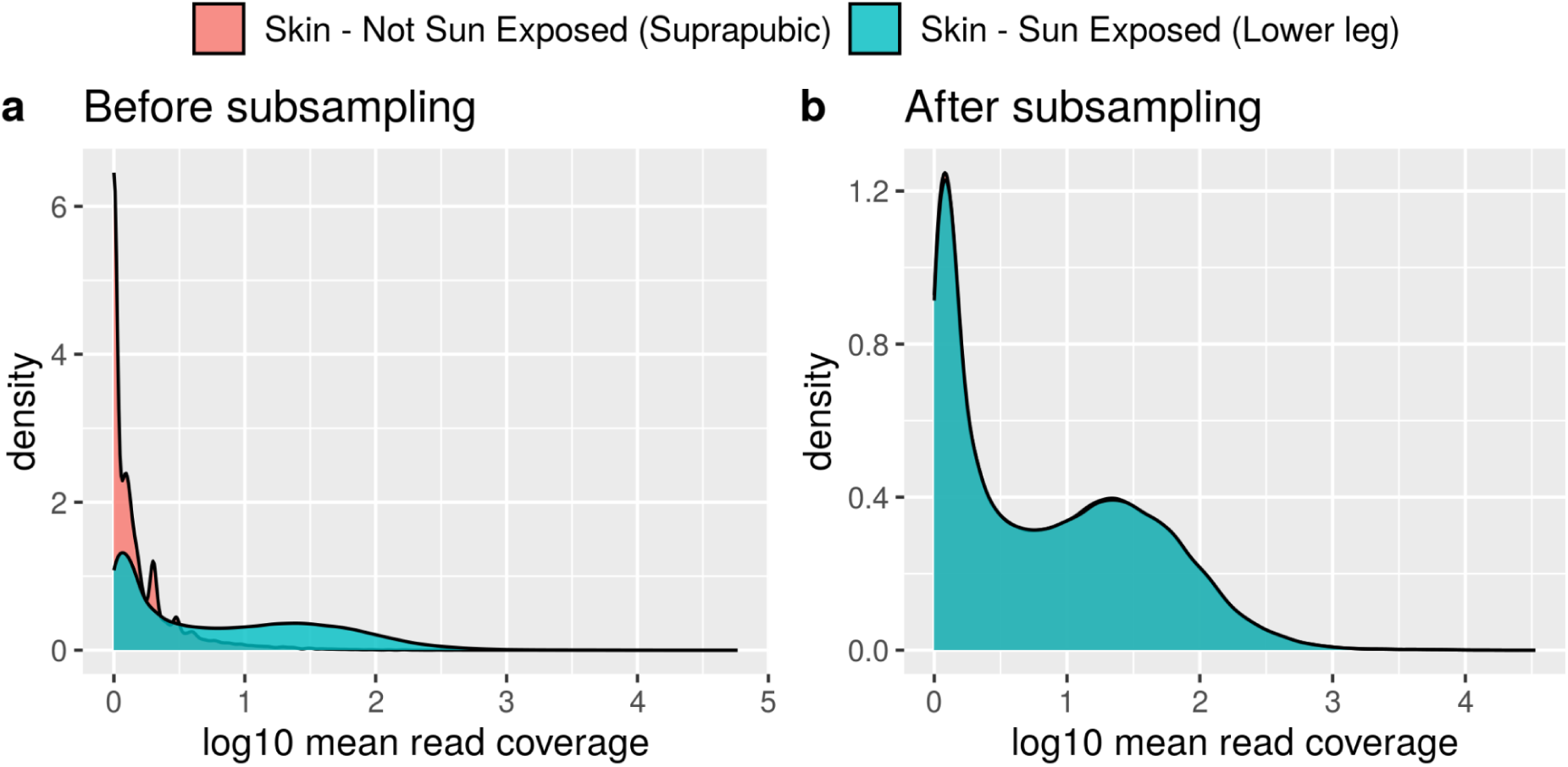
Mean read coverage of the annotated introns from “Skin non-sun-exposed” and “Skin sun-exposed” tissue before and after subsampling them by mean read coverage. **a.** Mean read coverage of the annotated introns from “Skin non-sun-exposed” tissue versus the annotated introns from “Skin sun-exposed” tissue before subsampling both distributions to meet for mean read coverage similarity. Only annotated introns showing a maximum difference of 0.005 in their mean read coverage between the two tissues were paired and kept for downstream analyses. **b.** Mean read coverage of the annotated introns from “Skin non-sun-exposed” tissue versus the annotated introns from “Skin sun-exposed” tissue after subsampling and pairing them by mean read coverage similarity. Only annotated introns showing a maximum difference of 0.005 in their mean read coverage between the two tissues were paired and kept for downstream analyses.

**Extended Data Fig. 6:**
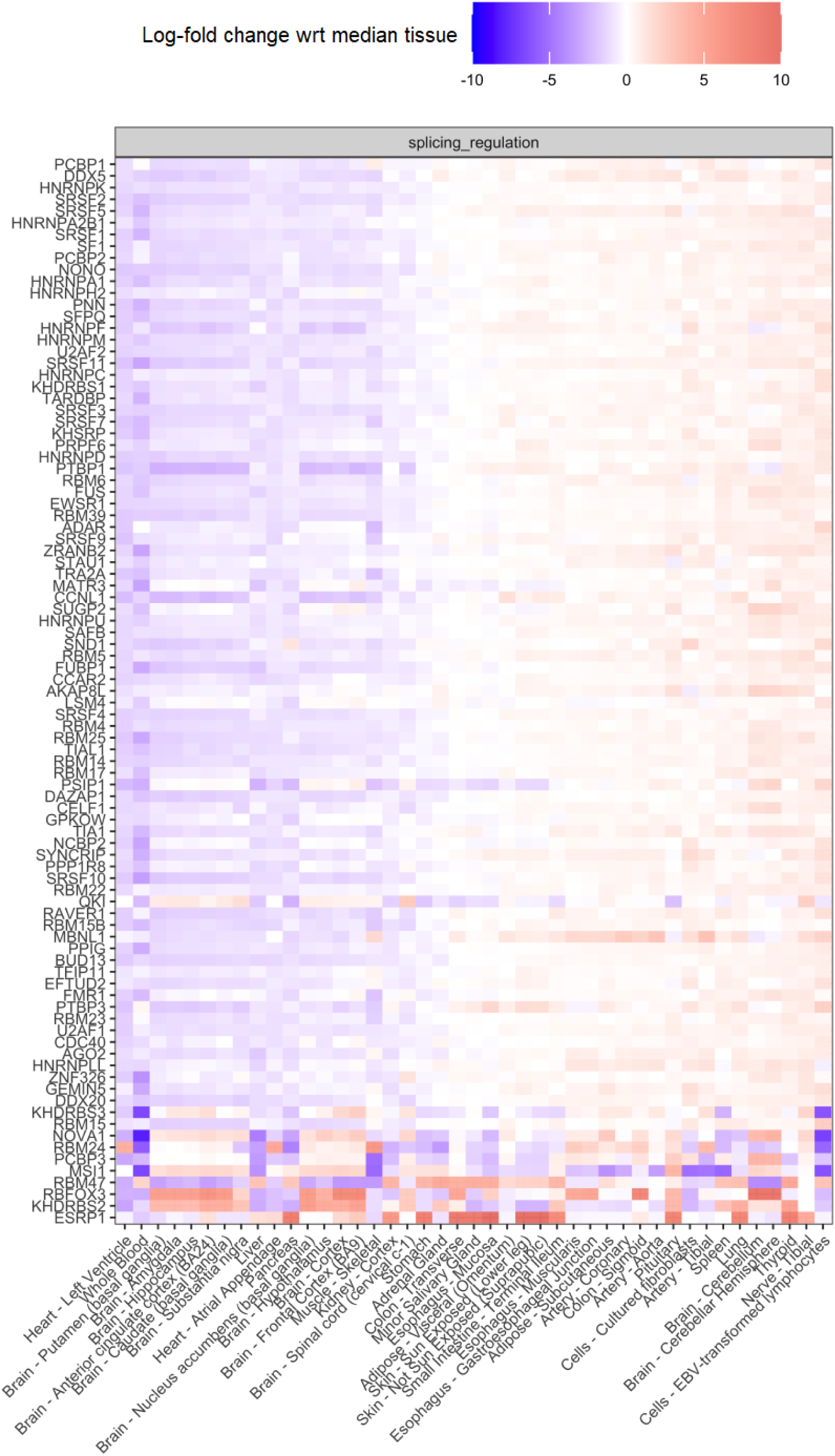
Median expression level of 98 RBPs in 42 GTEx body sites. Log fold-change median expression level of 98 splicing-regulator RBPs defined by Van Nostrand et al. 2020 across the samples of each one of the 42 GTEx body sites studied.

**Extended Data Fig. 7:**
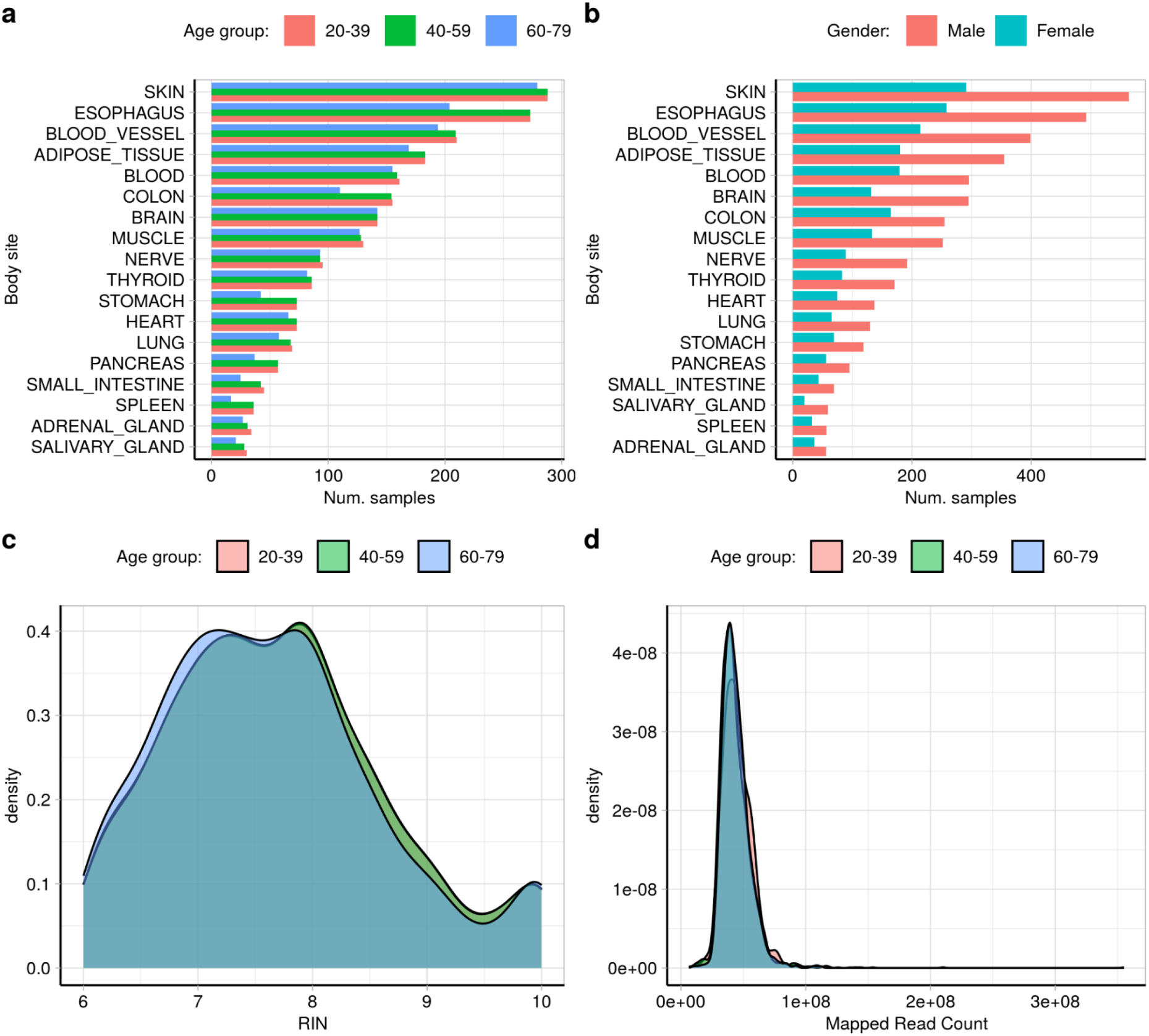
Metadata of the samples included in the “Age Stratification” intron database. All samples considered had been selected after subsampling and balancing them to meet by RIN number similarity across the three age groups.

**Extended Data Fig. 8:**
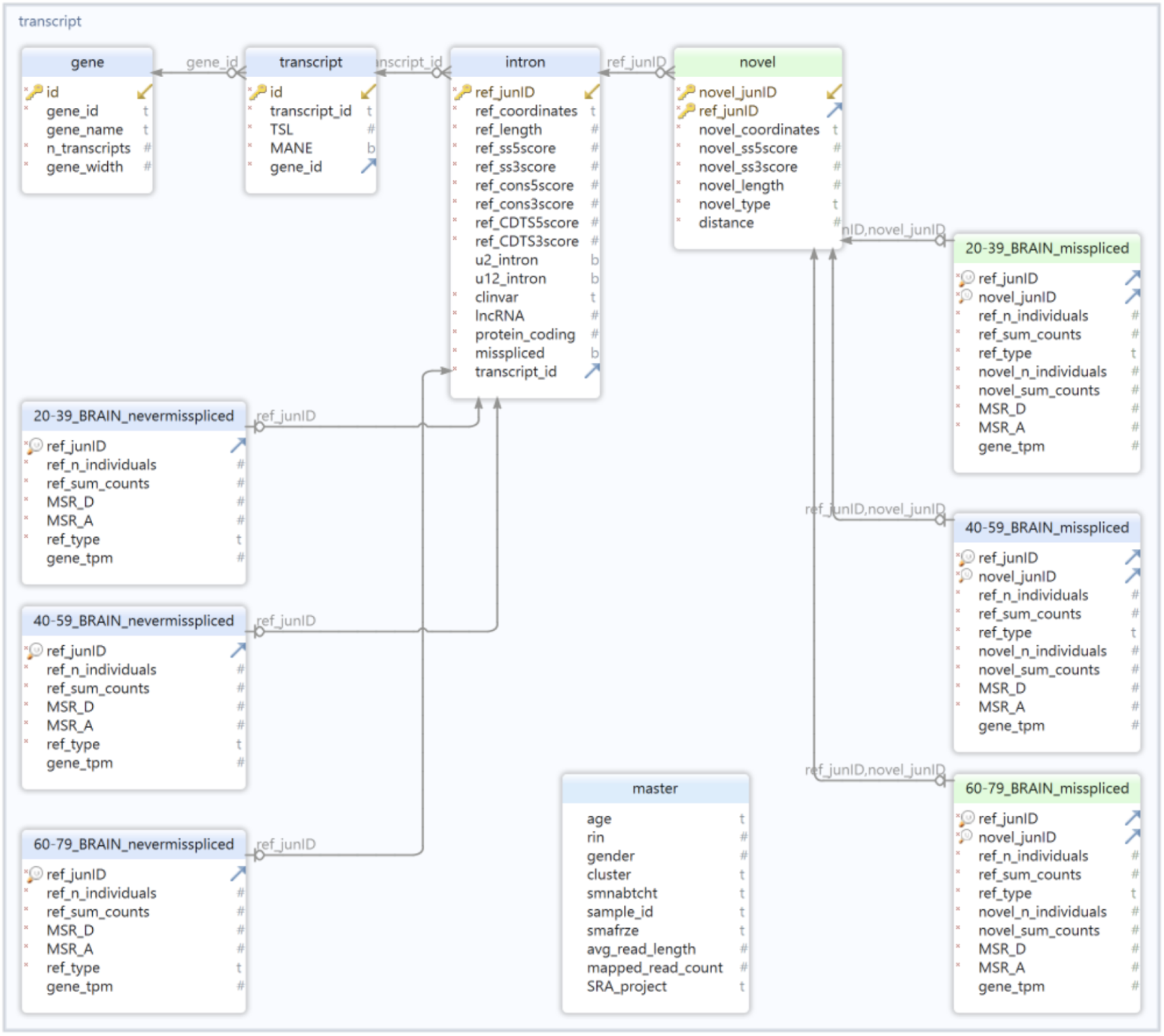
SQL schema of the “Age Stratification” intron database. To facilitate the visualisation of the database structure, only tables from the brain body site are shown.

**Extended Data Fig. 9:**
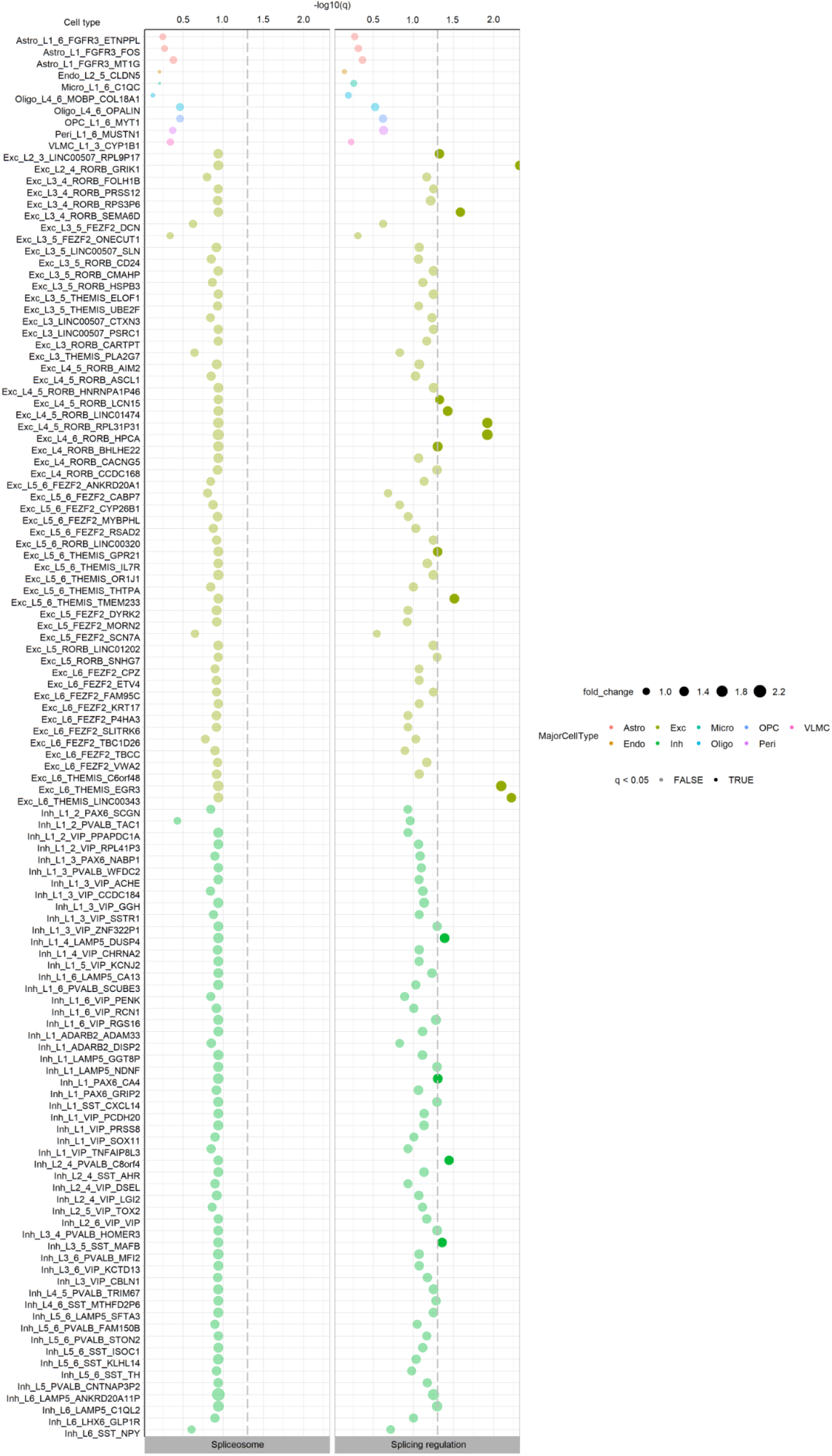
Cell-type specific expression of 98 splicing-regulator and 35 spliceosomal RBPs, defined by Van Nostrand et al. 2020, in multiple cortical regions from the human brain. The cell type annotations used correspond to the original clusters defined by the Allen Brain Atlas (Shen et al. 2012). The dashed grey vertical lines represent the minimum level of significance, with dots displayed on the right of the dashed line showing a significant expression for a given cell type. P-values were corrected for multiple testing using the Benjamini-Hochberg method, resulting in q-values.

**Extended Data Fig. 10:**
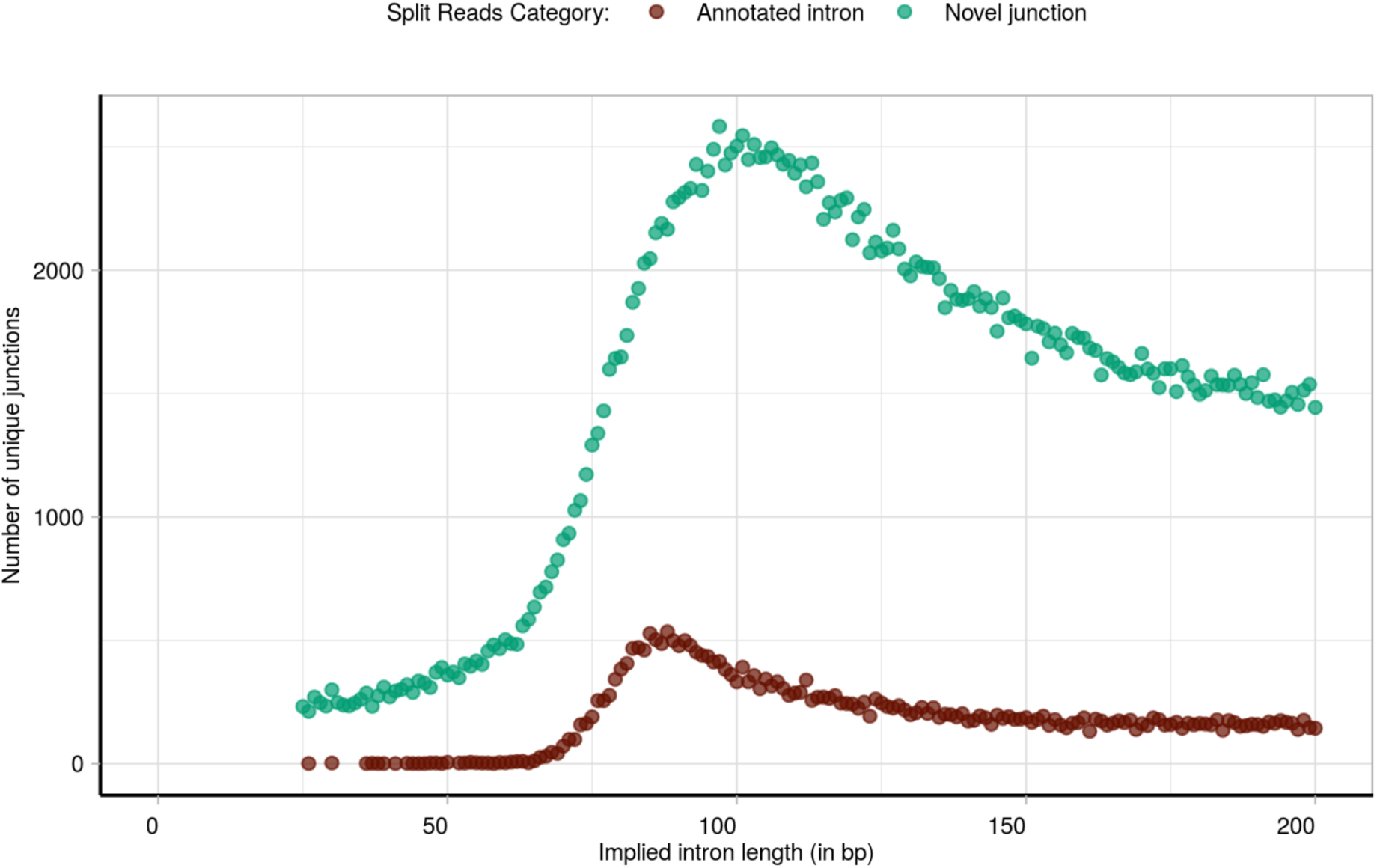
Overview of the implied intron length of the dataset of annotated and novel split reads studied. Distribution of the implied intron length corresponding to all novel donor and novel acceptor split reads studied (represented in green) compared to the implied intron length of the annotated split reads (in brown).

## Supplementary Figures

**Supplementary Fig. 1:**
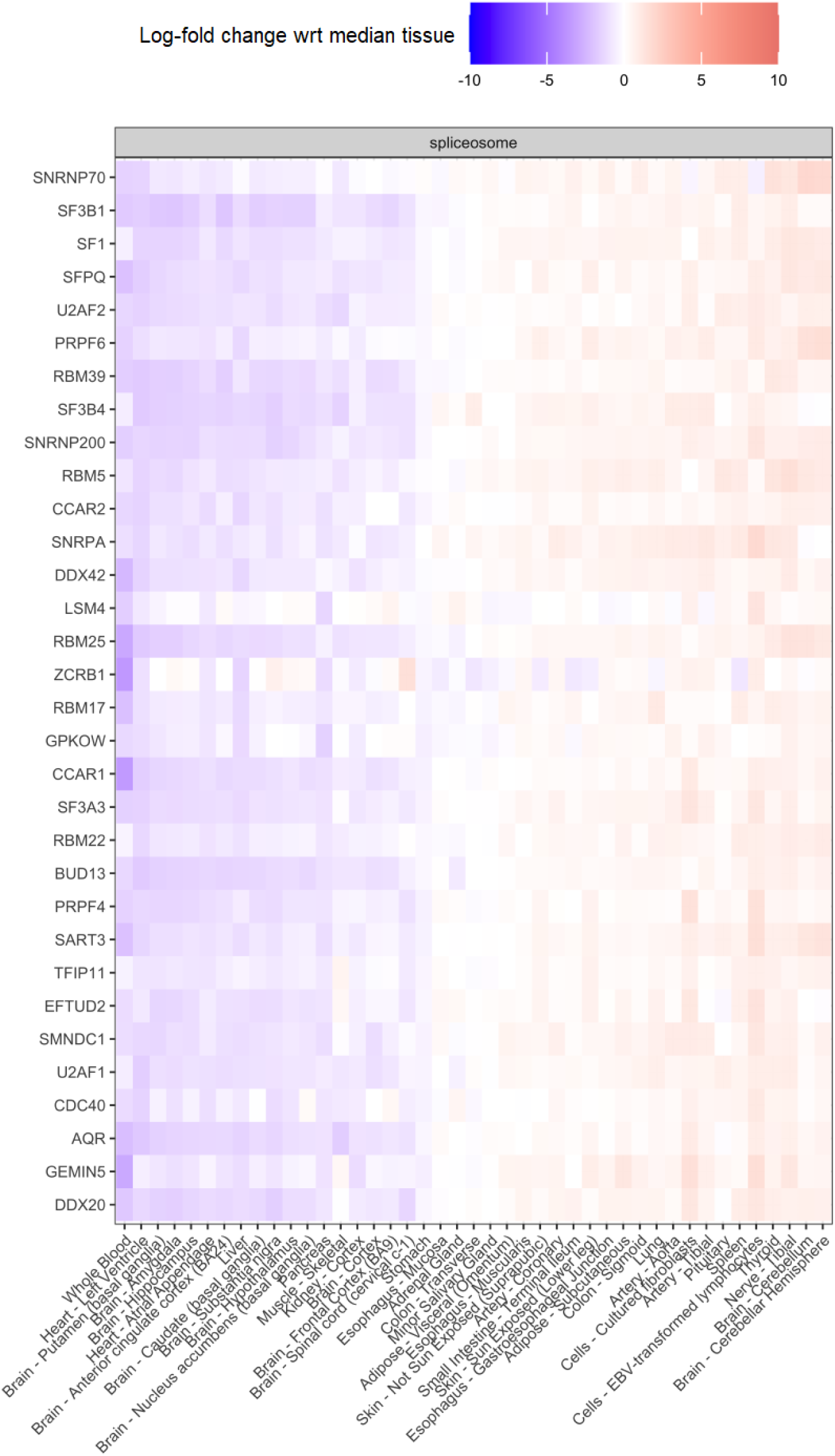
Median expression level of 35 RBPs in 42 GTEx body sites. Log fold-change median expression level of 35 spliceosome RBPs defined by Van Nostrand et al. 2020 across the samples of each one of the 42 GTEx body sites studied.

**Supplementary Fig. 2:**
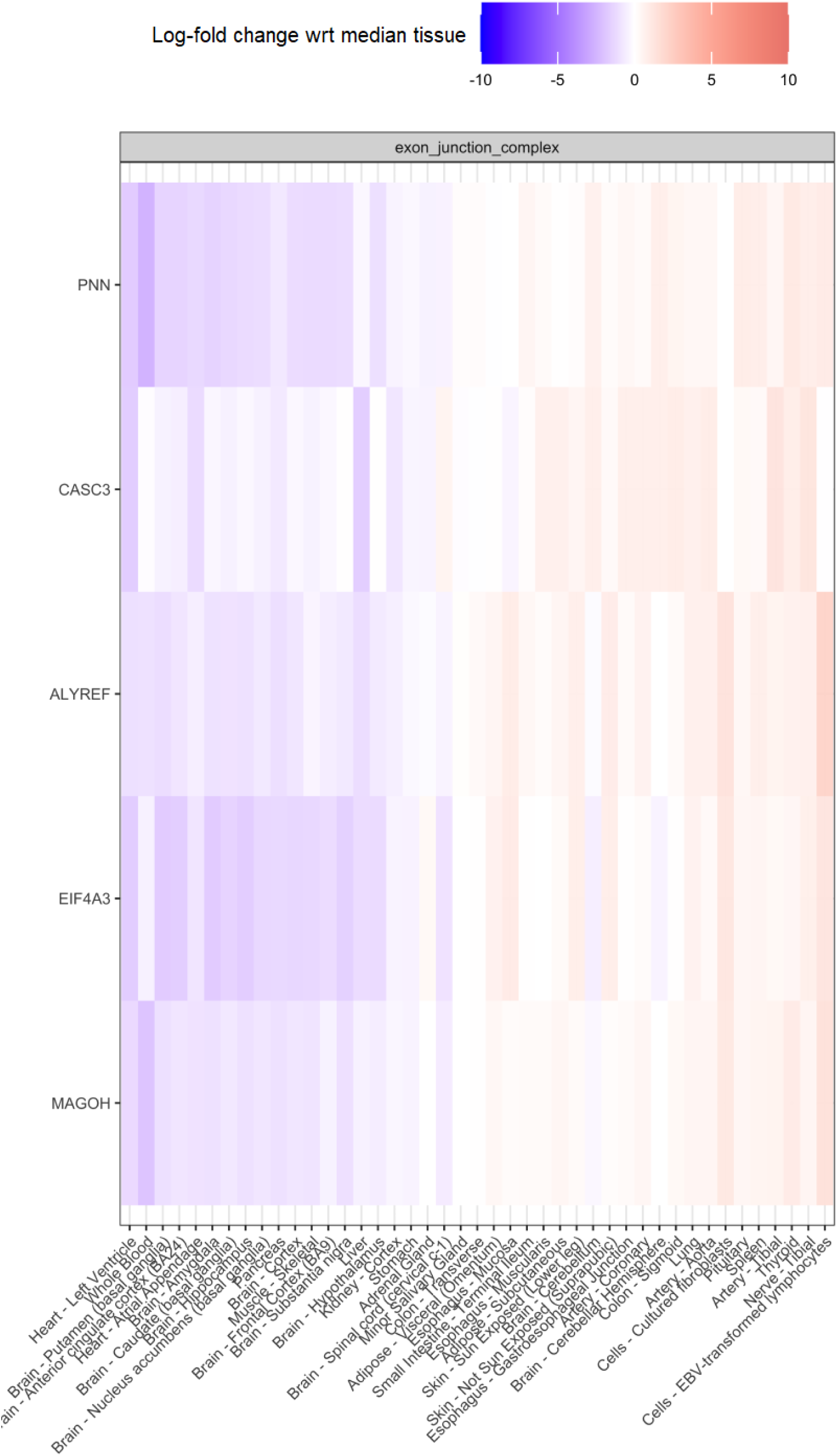
Median expression level of 5 RBPs in 42 GTEx body sites. Log fold-change median expression level of 5 exon-junction complex RBPs defined by Van Nostrand et al. 2020 across the samples of each one of the 42 GTEx body sites studied.

## Abbreviations

5’ss: Donor splice site
3’ss: Acceptor slice site
AGEZ: AG Exclusion Zone
bp: Base pair
BP: Branch Point
CDTS: Context-dependent tolerance score. It represents a measure of DNA sequence constraint in humans^66^
effsize: Probability of superior outcome between two compared groups^101^. It represents the probability that a randomly selected observation from group A will have a higher score than a randomly selected observation from group B. Unlike p-values, effect sizes are independent of the sample size
FCTX: Frontal cortex brain tissue
GO: Gene Ontology enrichment analysis
GTEx v8: Genotype-Tissue Expression (GTEx) v8 project^44^ (https://gtexportal.org/home/tissueSummaryPage)
IQR: Interquartile range
KEGG: Kyoto Encyclopedia of Genes and Genomes
MES: Maximum Entropy Scan score, http://hollywood.mit.edu/burgelab/maxent/Xmaxentscan_scoreseq.html
mod3: Modulo3 of a distance in base pairs. The modulo3 was calculated by dividing a distance figure in base pairs by 3 and obtaining the remainder of this division. mod3=0 reflects that the division by 3 has been exact; mod3=1; reflects that the division by 3 has produced value 1 as remainder; mod3=2 reflects that the division by 3 has produced value 2 as remainder
mRNA: Messenger RNA
*MSR_D_*: Mis-splicing Ratio calculated at the 5’ss (donor splice site) of a given annotated intron
*MSR_A_*: Mis-splicing Ratio calculated at the 3’ss (acceptor splice site) of a given annotated intron
NMD: Nonsense-mediated decay pathway
phastCons20: The mean interspecies conservation score across for 20 alignments (human, 16 primates, dog, mouse and tree shrew) to the human genome of the proximal intronic sequences (−5/+35bp, −35/+5bp, ‘/’ meaning exon-intron junction) tested. http://hgdownload.cse.ucsc.edu/goldenPath/hg38/phastCons20way/
PPT: Polypyrimidine Tract
pre-mRNA: Messenger RNA precursor
q: FDR-adjusted p-value. The False Discovery Rate (FDR) multiple testing adjustment method^102^ was formally described by Yoav Benjamini and Yosef Hochberg (i.e. Benjamini-Hochberg method) in 1995
RBP: RNA-binding Protein
shRNA: Short Hairpin RNA
TPM: Transcripts Per Million
V: V-statistic produced by a paired Wilcoxon rank sum test with continuity correction.

## Declarations

This manuscript used anonymised human RNA-sequencing data provided by the Genotype-Tissue Expression (GTEx) v8 project^44^ and processed by the recount3^83^.

## Acknowledgements

This research was funded in whole or in part by Aligning Science Across Parkinson’s [Grant numbers: ASAP-000478, ASAP-000509, and ASAP-000486] through the Michael J. Fox Foundation for Parkinson’s Research (MJFF). For the purpose of open access, the author has applied a CC BY public copyright licence to all Author Accepted Manuscripts arising from this submission.

S.G.R. and M.R. was supported through the award of a Tenure Track Clinician Scientist Fellowship (MR/N008324/1). E.K.G. was supported by the Postdoctoral Fellowship Program in Alzheimer’s Disease Research from the BrightFocus Foundation (Award Number: A2021009F). A.F.-B. was supported through the award of a Biotechnology and Biological Sciences Research Council (BBSRC UK) London Interdisciplinary Doctoral Fellowship;

Z.C. was supported by a clinical research fellowship from the Leonard Wolfson Foundation; A.L.G.-M. was supported by Fundación Séneca [21230/PD/19]; J.B. was supported through the Science and Technology Agency, Séneca Foundation, CARM, Spain (research project 00007/COVI/20);

L.C.-T. was supported by the National Institutes of Health (United States) [R01MH123567]. D.C.R. was supported by the Michael J. Fox Foundation - ASAP program (ASAP-000486).

